# Structural analysis of cholesterol binding and sterol selectivity by ABCG5/G8

**DOI:** 10.1101/2022.05.18.492512

**Authors:** Danny Farhat, Fatemeh Rezaei, Milica Ristovski, Yidai Yang, Albert Stancescu, Lucia Dzimkova, Sabrina Samnani, Jean-François Couture, Jyh-Yeuan Lee

**Affiliations:** Department of Biochemistry, Microbiology and Immunology, Faculty of Medicine, University of Ottawa, Ottawa, Ontario, Canada; Translational and Molecular Medicine Program, Faculty of Medicine, University of Ottawa, Ontario, Ottawa, Canada; Biochemistry Program, Faculty of Health Sciences, McMaster University, Hamilton, Ontario, Canada; Biochemistry Program, Faculty of Science, University of Ottawa, Ottawa, Ontario, Canada

**Keywords:** ABCG5, ABCG8, ABCG1, ABCG sterol transporters, cholesterol, sterol selectivity, X-ray crystallography, molecular docking

## Abstract

The ATP-binding cassette (ABC) sterol transporters are responsible for maintaining cholesterol homeostasis in mammals by participating in reverse cholesterol transport (RCT) or transintestinal cholesterol efflux (TICE). The heterodimeric ABCG5/G8 carries out selective sterol excretion, preventing the abnormal accumulation of plant sterols in human bodies, while homodimeric ABCG1 contributes to the biogenesis and metabolism of high-density lipoproteins. A sterol-binding site on ABCG5/G8 was proposed at the interface of the transmembrane domain and the core of lipid bilayers. In this study, we have determined the crystal structure of ABCG5/G8 in a cholesterol-bound state. The structure combined with amino acid sequence analysis shows that in the proximity of the sterol-binding site, a highly conserved phenylalanine array supports functional implications for ABCG cholesterol/sterol transporters. Lastly, *in silico* docking analysis of cholesterol and stigmasterol (a plant sterol) suggests sterol-binding selectivity on ABCG5/G8, but not ABCG1. Together, our results provide a structural basis for cholesterol binding on ABCG5/G8 and the sterol selectivity by ABCG transporters.

## 1. INTRODUCTION

Lipid homeostasis in mammalian cells is a critical process regulated in part by the ATP binding cassette (ABC) lipid transporters, particularly the ABCG and ABCA cholesterol transporters, such as ABCG1, ABCG4, ABCG5/G8 and ABCA1. As a critical component of cellular membranes, cholesterol comprises >50% of total cellular lipid content and is further utilized as the precursor molecule for steroid hormones that modulate gene regulation as well as bile acids that are required for nutrient absorption. To maintain cholesterol balance, excess cholesterol must be eliminated from cells and tissues via the reverse cholesterol transport (RCT) pathway by circulating high-density lipoproteins [1–3]. Within the human ABCG protein subfamily, all members function as lipid floppases, with the notable exception of ABCG2, that opts for a more relaxed approach to substrate efflux by working as the sole multidrug transporter (MDR) of the subfamily [4, 5]. In addition, the PDR subfamily of full-length transporters (ABCG2 homologues in *Saccharomyces cerevisiae*) are related to ABCG and sharing the structural fold and inverted topology. Moreover, ABCG transporters are becoming increasingly important with new findings uncovering their physiological roles in severe chronic diseases, notably such as cancer, atherosclerosis, diabetes and Alzheimer’s disease [6–11]. While these proteins have gained more notoriety, the efflux mechanisms have remained a mystery, with various proposed hypotheses that are yet to be agreed upon.

In mammals, ABCG lipid transporters consist of the heterodimeric ABCG5/G8 and the homodimeric ABCG1 or ABCG4 complexes. ABCG5/G8 is localized on the canalicular membranes of the bile ducts in the liver and the brush border of enterocytes in the small intestines, where it mediates the final step of the RCT [9,12–14]. ABCG1 and ABCG4 translocate cholesterol within the plasma membrane and in endosomes [15, 16] and play a key role in the regulating cholesterol balance in the lung, the brain and in macrophage-rich tissues [17–20]. ABCG5/G8 undergoes obligatory heterodimerization and is unique in its capability of preferential efflux for dietary plant sterols over cholesterol [8,9,21,22]. The mechanisms that govern such substrate selectivity within this protein subfamily, however, remains elusive. Based on the crystal structure of ABCG5/G8, we can now theorize that such substrate specificity among ABCG transporters may be the result of the difference in hydrophobic valve composition between the multidrug transporter: ABCG2 and the lipid transporters: ABCG1, ABCG4, ABCG5/G8 [23, 24]. In addition, a sterol-binding site was postulated at the membrane-transporter interface based on the crystal structure of ABCG5/G8 [25], whereas studies in recent ABCG1 structures [26, 27] have suggested sterol binding at the cytoplasmic end of the transmembrane domain (TMD).

The general TMD topology is shared among ABCG subfamily proteins. Each half transporter consists of six transmembrane helices (TMH) and the re-entry helices that are part of the extracellular domains. A connecting helix bridges the TMD to the nucleotide-binding domain (NBD), forming an amphipathic interface on the cytoplasmic leaflet of the lipid bilayers. Meanwhile a triple-helical bundle brings the TMDs and the NBDs together to create a compact and short-stacked transporter conformation [23–25]. At the cellular level, ABC lipid transporters have been shown to alter the composition of their local membrane environment. Such revisions may come from the subduing of lipid rafts by redistributing the high cholesterol content towards the surrounding membrane regions [16, 28]. Using *in vivo* and *in vitro* functional assays, we have previously characterized a loss-of-function (LOF) mutation ABCG5_A540F_, a conserved residue near a putative sterol-binding site in ABCG5/G8 [25, 29]. These findings suggested the likelihood of an access site that allows the binding of cholesterol binding prior to its re-localization within the membrane-embedded portion of the sterol transporters.

One of the significant challenges in studying the cholesterol-transporter interaction is to accurately identify the sterol molecules as either the transport substrates or the intrinsic component of cell membranes. The latter can also play regulatory roles in membrane protein functions, such as ABCG2 [30] or G-protein coupled receptors [31]. Besides experimental approaches to obtain atomic models by crystallography or electron microscopy, *in silico* molecular docking can allow researchers to predict the ligand-binding sites on a protein by using available structural models and ligand libraries. The availability of either ABCG5/G8 or ABCG1 structural can now make it possible to provide a molecular framework and to allow further structural analyses by using *in silico* tools, such as molecular docking or molecular dynamics simulation.

In this study, we first carried out amino acid sequence analysis of ABCG transporters, highlighting key conserved aromatic residues arranged in a spatially conserved pattern on TMH2. We solved the crystal structure of ABCG5/G8 in complex with cholesterol. The structure shows that an orthogonal cholesterol molecule fitting horizontally in front of A540, a conserved ABCG5 residue at this orthogonal sterol-binding site. *In vitro* cholesterol-binding assay showed an inhibition of sterol binding activity by the mutant ABCG5_A540F_/G8. To determine the differential sterol binding activities on ABCG sterol transporters, blind molecular docking analysis was conducted on wild-type (WT) ABCG5/G8 or ABCG1 in the presence of cholesterol or stigmasterol, as well as selective mutants ABCG5_Y432F_, ABCG8_N564P_ and ABCG5_A540F_. We also analyzed the cholesterol-binding patterns on the ATP bound states by using both experimental and homology models, showing the sterols only accumulated on the surface of the extracellular domain, possibly an intermediate state during the ATP-coupled sterol-transport cycle. Together with previous *in vivo* and *in vitro* observations, our results provide a structural basis for sterol selectivity of ABCG lipid transporters and a visual component to the sterol-transport mechanism.

## 2. RESULTS

### 2.1 Conserved phenylalanine cluster on THM2 of ABCG transporters

Recent structural determination of ABCG transporters by cryo-electron microscopy [26,27,32,33] confirms the general molecular architecture as initially revealed by the crystal structure of an apo ABCG5/G8 (Lee et al., 2016). The TMH2 and TMH5 of both subunits are adjacent to one another, forming a dimer interface within the transmembrane domains. This interface has been hypothesized to be a sterol translocation pathway based on sterol localization as observed in the EM structures. However, to what extent this pathway contributes to sterol translocation remains unclear. In this study, we used multiple sequence alignment to analyze the amino acid sequence conservation in the TMH2 of human ABCG1/2/4/5/8, PDR5 of the yeast *Saccharomyces cerevisiae* [34], and White protein of *Drosophila melanogaster* [35, 36](**Fig. 1**). Among these seven ABCG transporters, White is a close homologue to ABCG1, while the drug resistant PDR5 is an ABC drug transporter and an ABCG2 homologue. Using ABCG1 as sequence reference and analyzing 300 mammalian orthologs, each ABCG member includes a highly conserved phenylalanine (or tyrosine in ABCG5) at position F1 (**Fig 1A**). The adjacent position F2 shows similar conservation of hydrophobic residues, albeit with a more relaxed sequence homology than F1 (**Fig 1A**). Position F3 reveals a conserved phenylalanine in ABCG1, ABCG4 and the N-terminal half of PDR5 (PDR5-1), although the C-terminal half of PDR5 (PDR5-2) has a conserved phenylalanine just one residue off from F3. Position F4 shows the last highly conserved phenylalanine found on TMH2 (**Fig 1A**). These conserved phenylalanine residues are visualized through a Logo plot (**Fig 1B**), indicating at least two phenylalanine residues being flanked between positions F1 and F4 on the hydrophobic residue-rich TMH2.

**Figure 1.**
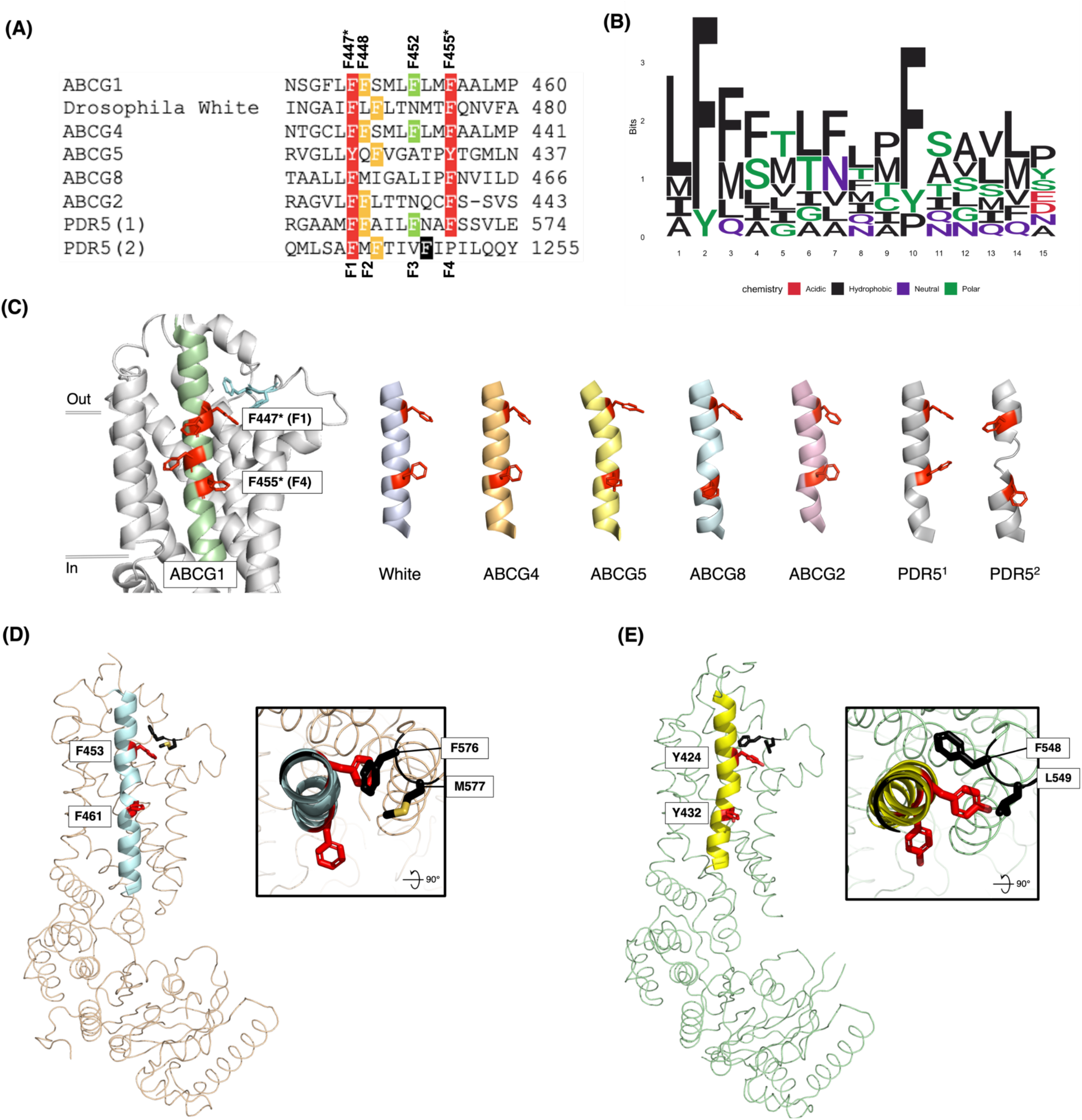
Conservation of aromatic amino acids on TMH2 of human ABCG family protein and homologues, including their localization in relation to key regions in the ABCG5/G8 heterodimer. **(A)** Sequence analysis of transmembrane helix five (TMH2) was done through PROMALS3D. Residues were screened for high conservation through BLAST sequence analysis. Red highlights perfect (or near perfect) aromatic residue conservation. Orange highlights conservation with ± 1 residue displacement. Green shows moderate conservation. Black shows non-sequence, but rather structural conservation. **(B)** Logo plot of TMH2 residues are colored according to their chemical nature. Red: acidic amino acids; Black: hydrophobic amino acids; Purple: neutral residues; Green: polar residues. **(C)** Structural conservation of highlighted aromatic amino acids from (A), using ABCG1 amino acid numeration as a reference. ABCG1 (PDB ID: 7OZ1, green), ABCG2 (PDB ID: 6VXI, pink, ABCG5/G8 (PDB ID: 5DO7; G5 in blue, G8 in yellow), PDR5-1 and 2 (PDB ID: 7PO3, light grey). *Drosophila* White (purple) and ABCG4 (orange) are the only two Alphafold models, both have helices with predicted local distance difference test score of >90. F447 (F1) and F455 (F4) residues in relation to ABCG1, have the highest conservation. The residues directions are maintained, with F447 stacking under the hydrophobic valve and F455 pointing towards the lumenal space between the TMD dimer. ABCG5 is the only monomer which has tyrosine’s instead of phenylalanine’s. ABCG1 and ABCG4 have large hydrophobic amino acids pointing lumenally causing a mass hydrophobic region spanning most of the transmembrane domain. **(D, E)** ABCG5/G8 heterodimer is split and shown in a top-down view, highlighting the location of conserved F453 and Y424 (red) in relation to hydrophobic valves (black). **(D)** In ABCG8, F453 (F1) stacks underneath F576, which (along M577) comprise the hydrophobic valve shown in black. **(E)** In ABCG5, Y424 (F1) is stacked underneath both hydrophobic valve residues; F548 and L549, shown in black.

When inspecting the spatial relationship of these phenylalanine residues, we observed an open space immediately below the hydrophobic valve and above a cavity that is open to the cytoplasm in the inward-facing conformation of ABCG transporters (**Fig. 1C, left**). Not only are the F1-4 residues shared among the eight homologous monomers, but two or three aromatic side chains also point inwards the TMD dimer interface, creating a highway-shaped open space (**Fig 1C, right**). We have coined this open space as the Phenylalanine Highway (PH) to highlight this unique structural motif in the transmembrane domain. ABCG1 and ABCG4 share the most closely spaced phenylalanine residues in their iteration of the PH, while the F2 in White is one residue off from that in ABCG1. ABCG2, ABCG8 and PDR5 (either the N- or C-terminal half) have similar spacing of their phenylalanine residues, whereas the equivalent residues on ABCG5 are replaced by aromatic tyrosine residues. In addition, the phenylalanine side chains may be adjusted based on the ligand placements (**Suppl Fig. 1**). Moreover, position F1 is spatially adjacent to the aromatic residues of the previously described hydrophobic valve (or ABCG2’s di-leucine valve/gate), forming a closed interface in the inward-facing protein conformation (**Fig 1D & 1E**) [23]. It may be possible to speculate that with such close spatial proximity, interaction between the hydrophobic valve and the phenylalanine residues could occur during the catalytic cycle via certain conformational changes. Further investigation is needed to address this question.

### 2.2 Crystal structure of ABCG5/G8 in an orthogonal cholesterol-binding state

Several *bona fide* sterol molecules were observed on recent cryo-EM structures of ABCG1 or ABCG5/G8, which led to various models of their sterol-transport (or binding) mechanisms, but more structural data and further mechanistic analysis are still required to establish a convergent model of cholesterol transport. We have previously crystallized human ABCG5/G8 in an apo state by bicelle crystallization. To gain the structural basis of the sterol-transport (or binding) mechanism, in this study, we determined a cytoplasmic-facing crystal structure of cholesterol-bound ABCG5/G8 at 4Å by X-ray crystallography. Three X-ray diffraction datasets were integrated and scaled with acceptable statistics. The initial model was obtained by molecular replacement using a previously resolved cryo-EM structure (PDB ID: 7JR7) [33], and the motifs involved in ATP binding were adjusted based on the previous crystal structure (PDB ID: 5DO7) [25]. Two ABCG5/G8 heterodimers are observed in the asymmetric unit. The protein model in this report includes a corrected amino acid registry in the nucleotide-binding domain as previously highlighted by Zhang *et al.* Additionally, it shows a consistent overall architecture and key structural features as described previously, overall root mean square deviation (RMSD) of ∼0.9 Å when superimposed [12,25,33](**Suppl. Fig. 1A**). After initial refinement, we observed two clear electron densities at the interface of symmetrically located transmembrane domains of these two transporters, which resembled the polycyclic characteristic of cholesterol molecules. Therefore, we added two cholesterol molecules to the initial model for further refinement and obtained a significant improvement of the Rfree from 0.338 to 0.302 and reasonable geometrical validation at current resolution. The final model shows an orthogonal cholesterol molecule in close contact with ABCG5_A540_ (namely the ABCG5-dominant side), with the 3-β-hydroxyl group forming a weak hydrogen bonding with the side chain of ABCG8_T430_ (4.3-4.5 Å). No sterol was observed on the equivalent side of ABCG8 (namely the ABCG8-dominant side) (**Fig. 2A**). Selected regions of the electron densities are shown in Suppl Fig. 2. The statistics of data processing and model refinement are summarized in **Table 1**.

**Figure 2.**
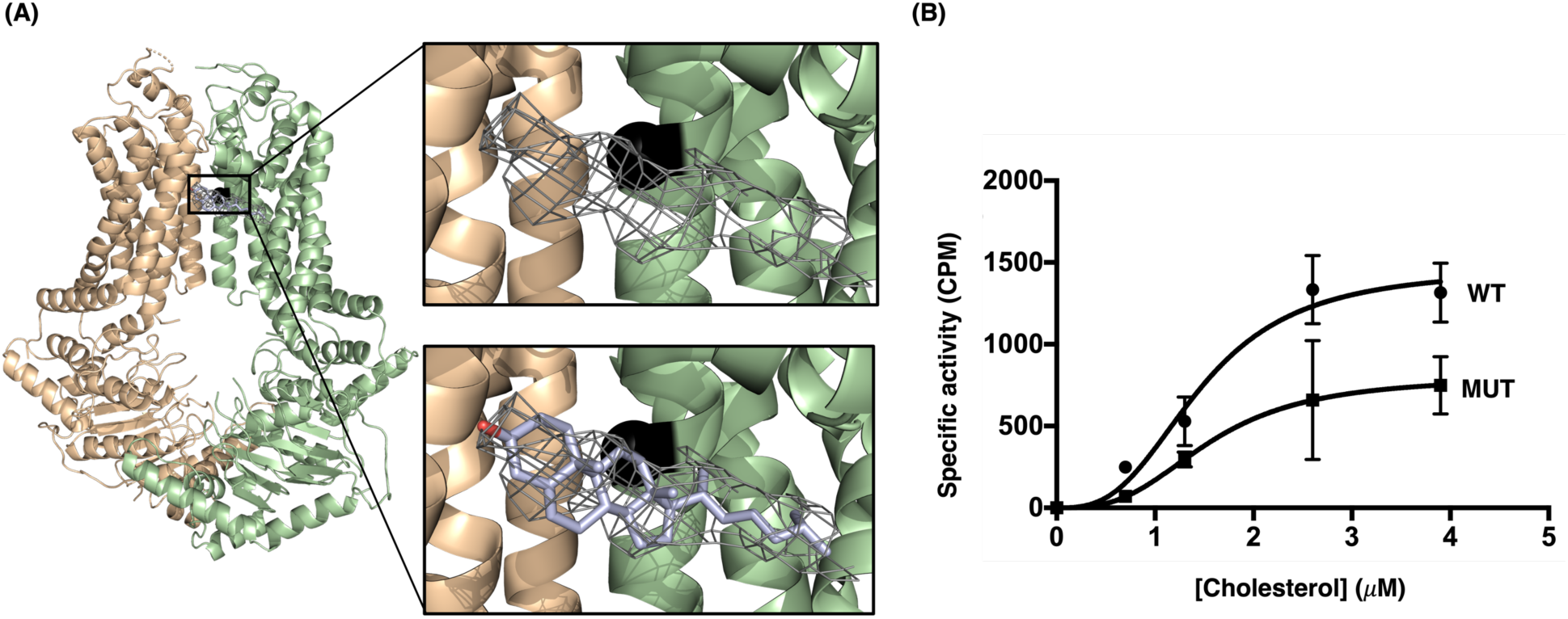
Crystal structure of cholesterol-bound ABCG5/G8 and cholesterol-binding assay. **(A)** Cholesterol-bound ABCG5/G8 structure is plotted in cartoon presentation. The first zoomed-in picture shows the electron density related to the cholesterol molecule which was observed at the interface of transmembrane domains of the crystallographic dimers (top right). The cholesterol molecule was modeled to the primary structure as shown in the second zoomed-in picture (bottom right). Alanine 540 (A540) is marked in a black sphere as a residue involved in the cholesterol-binding site. **(B)** The cholesterol-binding activities of the WT and the A540F mutant were measured by using [^3^H]-cholesterol, showing the binding parameters for the WT and the mutant with Bmax = 50.64±0.77 and 27.65±0.01 *μ*Ci/ml, K_D_ = 1.46 ±0.28 and 1.55±0.01 *μ*M, and h = 2.91 and 2.98, respectively. A P-value = 0.043 was obtained by paired t-test between WT and mutant. All calculations were performed using GraphPad Prism programs.

**Table 1:** Crystallographic data collection and refinement.

|  |  |
| --- | --- |
| **Beamline** | APS 19-ID |
| **Number of crystals** | 3 |
| **Wavelength** | 0.9794 |
| **Resolution range** | 30.58 - 4.051 (4.195 - 4.051)* |
| **Space group** | I 2 2 2 |
| **Unit cell (*a*, *b*, *c* (Å))** | 173.268 230.11 249.819 90 90 90 |
| **Total reflections** | 427824 (30232) |
| **Unique reflections** | 38173 (915) |
| **Multiplicity** | 11.2 (14.4) |
| **Completeness (%)** | 83.96 (22.75) |
| **Mean I/sigma(I)** | 5.90 (1.41) |
| **R-merge** | 3.392 (5.094) |
| **CC1/2** | 0.377 (0.137) |
| **CC\*** | 0.74 (0.491) |
| **Reflections used in refinement** | 34374 (915) |
| **Reflections used for R-free** | 2016 (54) |
| **R-work** | 0.2459 (0.2832) |
| **R-free** | 0.3029 (0.3039) |
| **CC(work)** | 0.864 (0.703) |
| **CC(free)** | 0.817 (0.715) |
| **Number of non-hydrogen atoms** | 18498 |
| **macromolecules** | 18442 |
| **ligands** | 56 |
| **RMS(bonds)** | 0.005 |
| **RMS(angles)** | 0.93 |
| **Ramachandran favored (%)** | 93.54 |
| **Ramachandran allowed (%)** | 5.90 |
| **Ramachandran outliers (%)** | 0.56 |
| **Rotamer outliers (%)** | 0.00 |
| **Clashscore** | 11.97 |
| **Average B-factor** | 52.45 |
| **macromolecules** | 52.45 |
| **ligands** | 49.48 |
| **Number of TLS groups** | 1 |

\* Statistics for the highest-resolution shell are shown in parentheses.

With an *in vivo* analysis of biliary cholesterol secretion and an *in vitro* ATPase assay, we have previously characterized a loss-of-function (LOF) mutant ABCG5_A540F_/G8 (A540F), predicting a putative sterol-binding at this transporter-membrane interface [25, 29]. In this study, we developed an *in vitro* cholesterol-binding assay using tritiated cholesterol (see Materials and Methods) and characterized cholesterol binding on purified ABCG5/G8 WT and A540F mutant proteins (**Fig. 2B**). The reaction reached a saturation of cholesterol at ∼3 μM with the B_MAX_ as 50.64±0.77 CPM, where the mutant showed about two-fold inhibition of the maximal cholesterol-binding rate. The binding constant K_D_ and Hill coefficient h were estimated as 1.46 ± 0.28 μM (h = 2.91) and 1.55 ± 0.01 μM (h = 2.98) for the WT and mutant proteins, respectively. Given the ATPase activity of A540F mutation was suppressed by ∼90% with similar K_M_ for sterols [29], both assays suggest a functionally defective conformation by this LOF mutant. Together with the new structural data, our analysis supports the role of the orthogonal cholesterol-binding site in proximity to ABCG5_A540_ (**Fig. 2**).

### 2.3 Asymmetric binding of cholesterol or stigmasterol on ABCG5/G8, but not ABCG1

The ABC transporter superfamily translocate a broad spectrum of substrates across cellular membranes, with individual members conveying their unique substrate specificities. To determine how ABCG transporters bind sterol ligands, we used previously published apo models ABCG5/G8 (PDB ID: 5DO7 or 7JR7) and ABCG1 (PDB ID: 7R8D) [25, 33] and carried out molecular docking on the TMD with cholesterol or stigmasterol, a plant sterol previously shown to have the highest accumulation rate in an Abcg5/g8-deficient mouse model [37]. Using UCSF DOCK6 suite [38], we first optimized the docking protocol by carrying out a redocking of the ligands (topotecan, mitoxantrone and imatinib) to reproduce their complex with ABCG2. Additional redocking was done on a distinct ABCG2 model to further evaluate docking optimization. (**Suppl Fig. 3**) [39, 40]. Figure 3 shows the docking results of all acceptable (or viable) poses for the two sterol ligands, cholesterol and stigmasterol, on the ABCG sterol transporters according to the standard flexible docking protocol as implemented in DOCK6 suites [38]. On ABCG5/G8, poses of sterols are distributed asymmetrically at the opposing membrane-transporter interfaces, having more docking conformations on the ABCG5-dominant side (**Fig. 3A/B**). The predicted orthogonal cholesterol (*i.e.*, one of the top-3 poses) has a nearly identical position to the cholesterol molecule in the newly determined cholesterol-bound structure (**Fig. 2 & 3D/E**). On the ABCG8-dominant side, no orthogonal sterol was obtained, and either ligand are clustered on the inner leaflet of the membrane’s lipid-bilayered and with the topmost sterols in close distance from the lower end of the PH (*i.e.*, ABCG5_Y432_ or ABCG1_F455_). Such asymmetric sterol binding on ABCG5/G8 suggests another level of functional asymmetry by ABCG5/G8-mediated sterol transport. On the ABCG1 homodimer, acceptable poses are only observed on the inner leaflet of the lipid-bilayered membranes, resembling those on the ABCG8-dominant side, and bound sterols can be equally distributed on either side of the transporter-membrane interface (**Fig. 3A/B, bottom**). Such symmetric sterol binding is distinct from ABCG5/G8, suggesting different mechanisms of using the PH between ABCG1 and ABCG5/G8.

**Figure 3.**
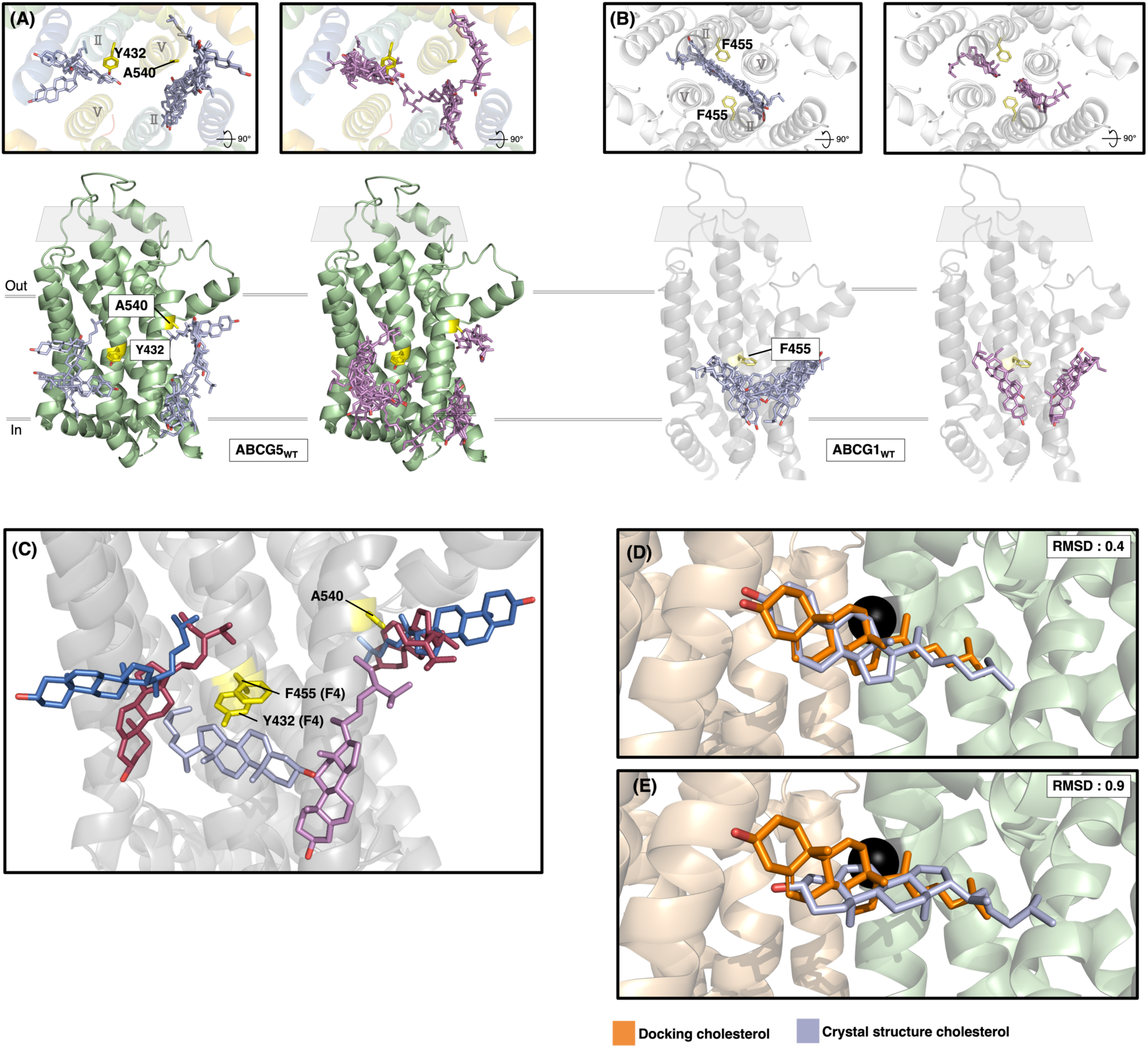
Cholesterol and stigmasterol docking in ABCG5/G8 and ABCG1. **(A-B)** Cholesterol (blue, left) and stigmasterol (violet, right) are bound to either side of the interface between the two dimers. TMH2 and TMH5 are labeled as II and V. **(A)** Cholesterol or stigmasterol in ABCG5/G8 docks differently on either side (shown only ABCG5 here). *Left of Y432*, cholesterol docks in three conformations: lightly angled upwards above Y432, horizontal on the X-axis below Y432 and vertical. Stigmasterol (right) poses to the left of Y432 has a “C” shaped pattern that surrounds Y432. *Right of Y432*, cholesterol binds in a continuous pattern, starting at A540 and ending at the lower edge of the transmembrane domain (TMD). A single cholesterol conformation binds horizontally near A540. Stigmasterol binds in two separated clusters with the top cluster showing horizontal poses ahead of A540 and the bottom cluster containing horizontally oriented sterols. **(B)** On ABCG1 (PDB ID: 7OZ1), either cholesterol or stigmasterol includes similar docking poses on both sides. The sterols are angled towards the putative sterol binding cavity on the cytoplasmic end of the bilayers. Given similar binding pattern, more cholesterol poses are obtained than stigmasterol. **(C)** Top sterol poses on both transporters are illustrated as a superposition. ABCG1 cholesterol and stigmasterol are colored blue and violet respectively. ABCG5/G8 cholesterol and stigmasterol are colored dark blue and dark red respectively. In general, sterols in ABCG5/G8 dock higher, near the orthogonal sterol-binding site, whereas sterols in ABCG1 binds under F455 (below F4 of the phenylalanine highway). **(D, E)** The cholesterol docking pose (orange) is aligned with either cholesterol in the asymmetric unit of the crystal structure (light blue) with an RMSD of 0.4 Å (D) and 0.9 Å (E).

To analyze sterol docking with ATP-bound structures, we performed the analysis on an ATP-bound ABCG1 and a homology model of ATP-bound ABCG5/G8 (see Materials and Methods) [41]. Surprisingly, given the same docking grid as used for apo structures, sterol docking on the ATP-bound ABCG1 or ABCG5/G8 only showed clusters on top of the extracellular domains of the extracellular-facing structures and far away from the PH (**Fig. 4A/B**). As current apo or ATP-bound structures of ABCG sterol transporters are rather limited to early and end states of catalysis [25–27,33], the sterol poses as observed here may represent the post-efflux conformations where the previously observed ligand binding sites have become inaccessible due to the compression of protein’s interfacial cavity in these ATP-bound structures. For this reason, binding sites within the protein were unable to be detected and subsequently docked. We additionally utilized MOLE2.5 [42] to determine the volume of cavities detected within the TMD. In ABCG1 cavities substantially large were found in between the transmembrane helices, however these were away from the central canal of the protein (**Suppl Fig. 5B**, left). In ABCG5/G8, while one big cavity appeared within the central canal of the protein, it was found to be a conglomeration of multiple spheroidal shaped cavities bridged together by very thin regions. Each cavity on its own was presented to be far too small of a volume at around ∼200Å^3^ comparing to the volume of cholesterol (**Suppl Fig. 5B**, Right), being ∼627Å^3^ [43]. Our data thus emphasizes the need for occluded turnover state models, as seen in ABCG2 models [44].

**Figure 4.**
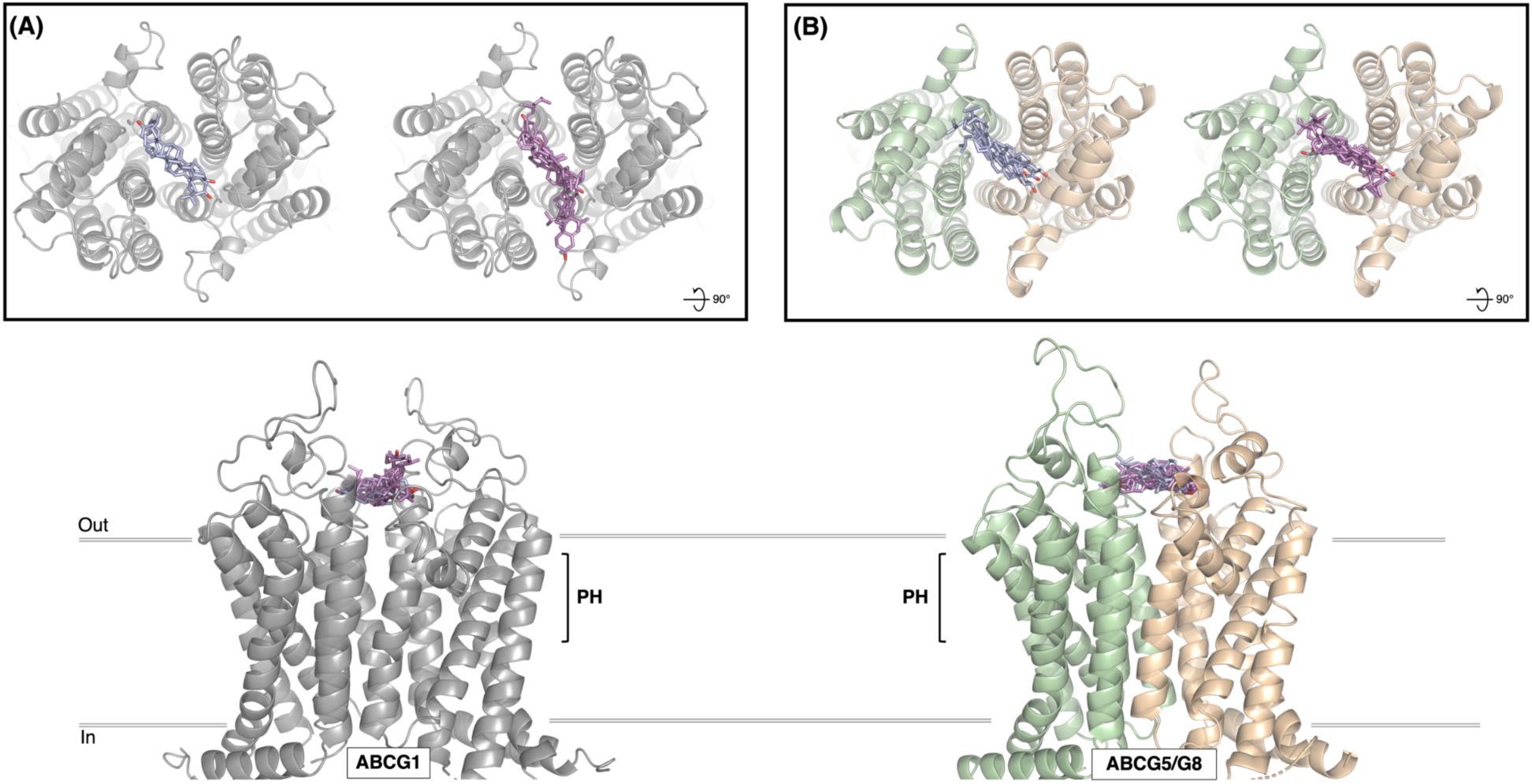
**(A, B)** Sterol docking on ATP-bound ABCG1 (A) and ABCG5/G8 (B) models. ABCG1 (PDB ID: 7R8E) in gray; ABCG5/G8 in pale-green and tan. Cholesterol (blue) and stigmasterol (violet) docked shown in a front view (lower) and a top-down view (upper). In brackets is the extent of the phenylalanine highway (PH) within the transmembrane domain (TMD) of the protein.

**Figure 5.**
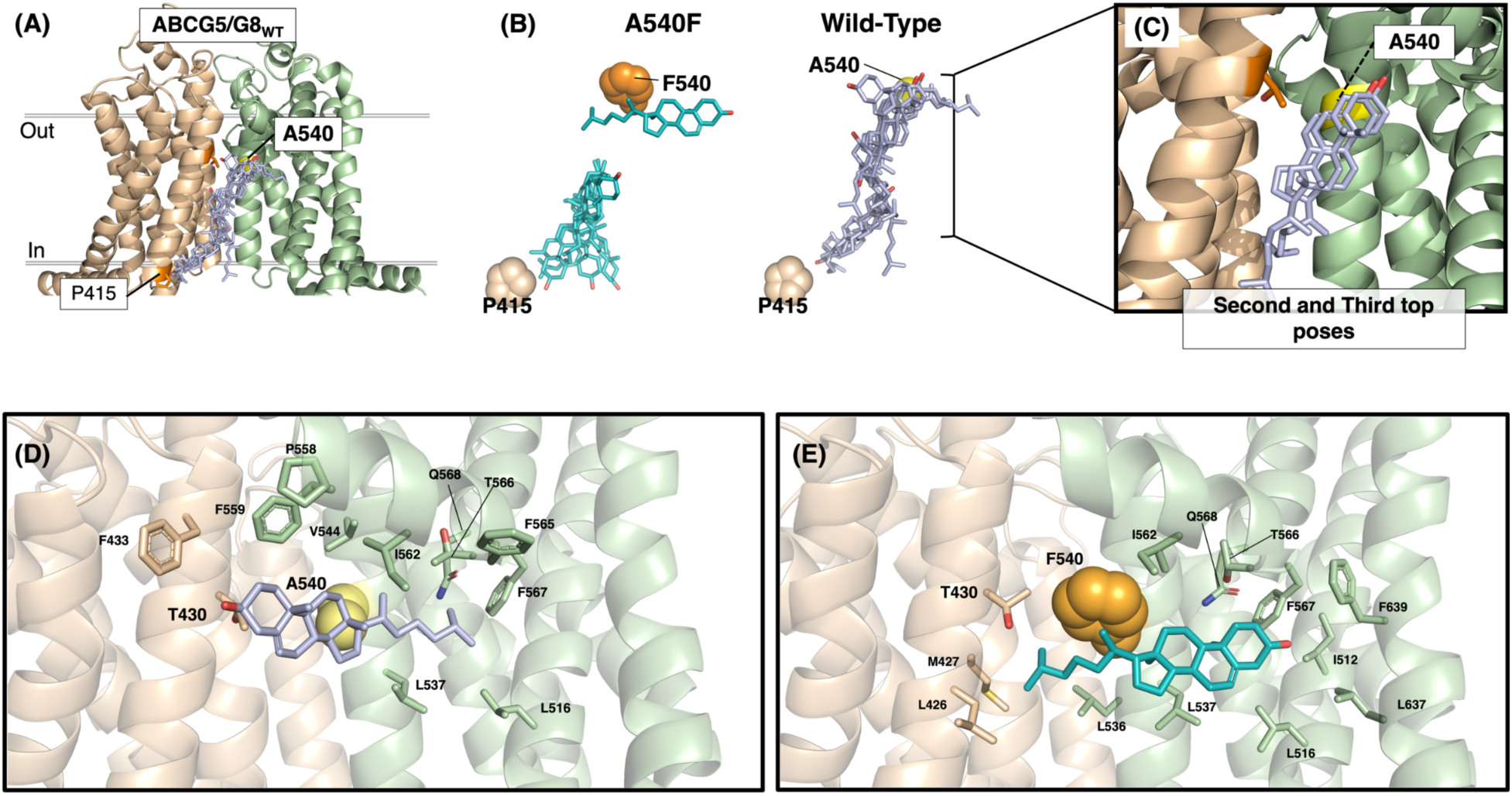
Wild-type and mutant A540F ABCG5/G8 docking predictions. **(A-C)** ABCG5 and ABCG8 are colored in pale-green and tan respectively, and the cholesterol docking on WT results are shown in (A). In comparison, WT docking is shown in blue while mutant is shown in teal (B). Landmark residues ABCG5_A540_ and ABCG8_P415_ are highlighted in tan and orange respectively for the orientation (B), showing that WT has a continuous cholesterol binding pattern from A540 to P415 while the mutant has a break in the docking prediction, separating the two clusters. Two of the top 3 poses in the wild type docking appear to be missing in the A540F mutant (C). **(D, E)** The micro-environment surrounding cholesterol highlights sterol-protein interaction. Residues are shown as sticks. From left to right, ABCG8: L426, M427, T430; ABCG5: F559, F540, L536, I562, L537 Q568, T566, F567, L516, L637. The side chain of F540 is shown in spheres to highlight the mutation.

### 2.4 Sterol selectivity by ABCG1 and ABCG5/G8

Among mammalian ABCG sterol transporters, ABCG5/G8 is known to promote absorption of cholesterol but not dietary sterols in the gastrointestinal tract [9]. To study such sterol selectivity, we analyzed the distributions of the sterol-binding poses from the docking experiments by using ABCG1 and ABCG5/G8 as model systems. On ABCG1, cholesterol cluster was gathered uninterruptedly around F455, *i.e.*, F4 of PH, and docked similarly on both sides of the transporter-membrane interface. The sterol cloud is angled towards the putative sterol binding cavity at the NBD-TMD interface, in the membranes lower leaflet. (**Fig. 3A, bottom**). Stigmasterol equally shows the same angling into the binding site, but with reduced possible poses when compared to cholesterol on ABCG1 (**Fig. 3B, bottom**). The largest number of positions for cholesterol or stigmasterol are located immediately below the PH and above the NBD-TMD interface, suggesting a common sterol-binding cavity in ABCG sterol transporters. However, the orientation the hydroxyl group on the sterols is not necessarily exposed to the aqueous milieu (**Fig. 3A/B, bottom**).

On the ABCG5-dominant side of ABCG5/G8, stigmasterol binds in two clusters partitioned between each monomer, leaving a gap in the middle, which is opposite to the continuous cloud as revealed by cholesterol (**Fig. 3A/B, top**). In general, there are more possible binding poses in the vicinity of ABCG5_A540_ and ABCG8’s PH, where interestingly all sterol molecules appear to orient in a similar direction, *i.e.*, the hydroxyl group pointing to the NBD-TMD interface. On the other hand, on the ABCG8-dominant side and near F4 of ABCG5’s PH, the sterol molecules are oriented in various but seemingly unspecific directions (or conformations) (**Fig. 3A/B**). When compared to cholesterol which binds in two separate pockets (**Fig. 3A, top**), stigmasterol has three binding pockets across both sides of the transporter-membrane interfaces (**Fig. 3B, top**). Interestingly, the highest clusters of stigmasterol and cholesterol are located on the opposite sides of the transporter, suggesting preferred association of sterol species on either side of the transporter-membrane interface.

ABCG1_F455_ and ABCG5_Y432_ are at the equivalent position F4 of the PH. When comparing the sterol binding patterns around these key residues, more cholesterol poses can be found around ABCG1_F455_ in an increasingly linear shape, while more stigmasterol poses were observed around ABCG5_Y432_ in a “C” shape (**Fig. 3A/B**). In the nucleotide-free structures as used for analysis here, sterol binding in general is positioned more towards the orthogonal sterol-binding site in ABCG5/G8 than in ABCG1. Sterol poses in ABCG1 cluster closer to the central cavity, around PH’s F4, while sterols bind ABCG5/G8 closer to the transporter-membrane interface (**Fig 3C**). The sterols of the top binding poses all have hydroxyl groups pointing outwards and away from PH’s F4 position. Interestingly, the cholesterol molecules are similarly angled in both ABCG1 and ABCG5/G8, *i.e.,* in parallel to the lipid bilayers.

### 2.5 Alteration of ABCG5/G8 cholesterol-binding activity by the LOF mutant A540F

By exploiting the LOF mutant A540F, *in vivo* and *in vitro* functional studies support the notion that it is exposed to the orthogonal cholesterol-binding site [25, 29] (**Fig. 2**). Using molecular docking and homology modelling, it is possible to develop hypotheses for the structural basis of how this mutation impacts ABCG5/G8 function. In this study, we analyzed the effects of inducing the A540F mutation on the binding positions of cholesterol as compared to in WT ABCG5/G8 (**Fig. 5**). In the WT protein, the cholesterol cloud stretches from proline 415 (P415, end of the connecting helix of ABCG8 and a landmark position) to A540 (**Fig. 5A**), with one of the top poses being the orthogonal sterol adjacent to A540. The remaining top-three are vertically oriented, exposing the hydroxyl group to the orthogonal sterol-binding site (**Fig. 5B, right & Fig. 5C**). Using a homology model containing the A540F mutation, the second and third best binding positions were missing, leaving a gap between PH’s F4 and the orthogonal sterol-binding site (**Fig. 5B, left & 5C**). As mutation was induced, the orientation of the orthogonal cholesterol is flipped 180° at the sterol-binding site (**Fig. 5D/E**), and the remaining non-top three poses have their hydroxyl groups pointing to the cytoplasmic side of the lipid bilayer (**Fig. 5B**). Overall, lower number of different cholesterol poses were docked on the mutant as compared to the WT protein. The microenvironment surrounding the orthogonal cholesterol on A540F mutant are highlighted in **Figure 5E**, showing that the phenylalanine substitution prevents the interactions between the cholesterol and polar residues (*e.g.*, ABCG8_T430_) that may attract the hydroxyl group in the WT protein.

### 2.6 Lower transmembrane sterol-binding cavity in ABCG5/G8

The recent cryo-EM structural determination of ABCG1 suggested a cholesterol-binding cavity in the cytoplasmic leaflet of the bilayer [26, 27]. The protein’s surface within the binding cavity in ABCG1 shows a narrow gap that is wider than corresponding cavity found in ABCG5/G8 (**Fig. 6A**); additionally, this cavity is located immediately below F4 of the PH (**Fig. 6B**). In the WT proteins, ABCG5_Y432_ and ABCG8_N564_ are separated by 4.0 Å, whereas the equivalent positions in ABCG1_F455_ and ABCG1_P558_ are farther separated by 6.9Å. We created an ABCG5/G8 homology model that consists of mutations ABCG5_Y432F_ and ABCG8_N564P_ to mimic an ABCG1-like cavity. The interaction between the mutant residues in ABCG5/G8 are at a similar distance and positions to the corresponding residues in ABCG1 (**Fig. 6B/C**). This distance is significantly widened from 4.0 Å to 6.9 Å in the mutant. Due to the increased distance between the residues, the double mutant lacks the ability to form such N-Y blockade and results in a deeper cavity than that of WT. We then performed sterol docking analysis on this double-mutant model and analyzed the possible sterol conformations in this cytoplasmic sterol-binding cavity. In the WT, stigmasterol appears to bind more readily than cholesterol, where the cluster of docked stigmasterol poses is much bigger than the cholesterol cloud (**Fig. 6D/E**). Both sterol clouds semi-encircle Y432 in a “C” shape. When Y432 is mutated to phenylalanine, the number of docked sterols decreased significantly as compared to the WT, simply localizing overhead of position 432 and suggesting lower sterol-binding ability in the mutant. Interestingly, the size of viable poses becomes more alike for either cholesterol or stigmasterol. Despite using ABCG5_Y432F_/G8_N564P_ mutations to mimic the width of the cavity found in WT ABCG1, the binding pattern in the mutant does not resemble that of ABCG1 (**Fig. 2A and Fig. 2B, bottom**), suggesting the possibility of either sterol transporter adopting multiple conformations to bind sterols throughout the transport cycle. Further functional assays are necessary to define the role of these residues and how they can obstruct sterol translocation trough the cavity.

**Figure 6.**
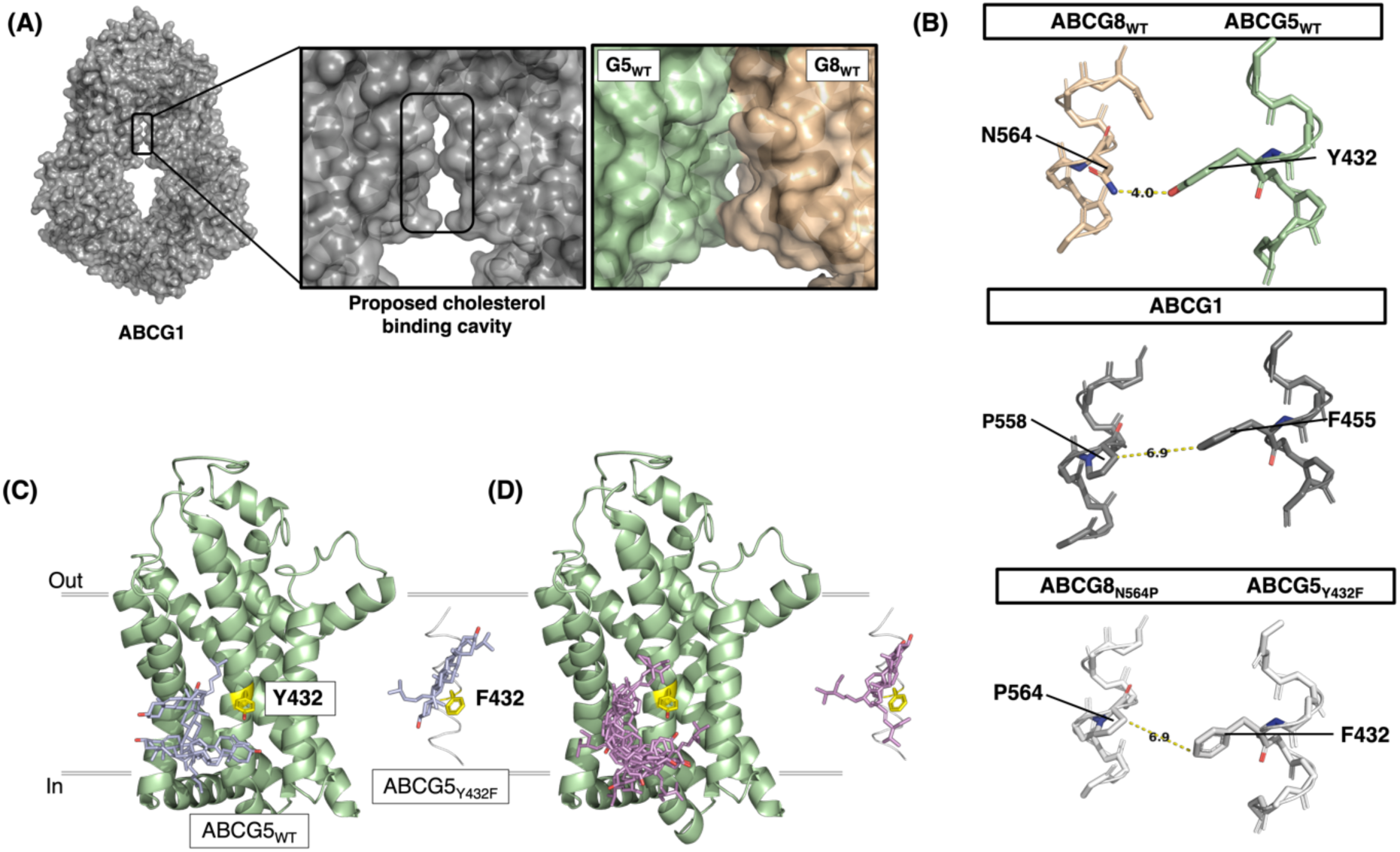
Lower transmembrane cavity in ABCG5/G8 and ABCG1 with cholesterol and stigmasterol docking on wild-type and double mutant (Y432F, N564P) ABCG5/G8. **(A)** ABCG1’s (PDB ID: 7OZ1, gray) hollow transmembrane region was previously denoted as a putative cholesterol binding cavity contrasted with ABCG5 (pale-green)/G8(tan). All atoms are included (but not displayed) with the surface fit over the protein. **(B)** WT ABCG5/G8 (PDB ID: 7RJ7, pale-green/tan) with residues Y432 and N564 shown as sticks. Distance between the two amino acids is ∼4.0 Å, and for the equivalent residues in ABCG1 (PDB ID: 7OZ1, gray), F455 and P558, they are ∼6.9 Å apart. A homology model of the double-mutant ABCG5/G8, shown in white, has Y432 and N564 substituted for phenylalanine and proline, respectively, as observed in ABCG1. **(C, D)** Cholesterol (blue) and stigmasterol (violet) docking was simulated on WT and double mutant (white) ABCG5/G8, with only ABCG5 shown here. Yellow: residue 432 on ABCG5. Wild type has more potential binding poses than the double mutant that only binds a handful sterols above residue 432.

## 3. DISCUSSION

Atomic models of ABCG sterol transporters have provided the molecular framework to study how individual transporter proteins translocate different substrates. Yet, the location and methods by which these transporters bind sterols remains elusive and controversial. This is particularly true in the case of ABCG5/G8, which is believed to identify and bind plant sterols differently from cholesterol. Understanding how sterols binds on ABCG5/G8 and its homologs will allow us to study the impact of such sterol selectivity on the regulation of sterol transport. In this study, we have determined a cholesterol-bound crystal structure of ABCG5/G8, showing an orthogonal cholesterol binding on the ABCG5-dominant side at the transporter-membrane interface (**Fig. 2**). A mutant ABCG5_A540F_ was previously demonstrated to downregulate biliary cholesterol efflux *in vivo* [25] and to inhibit sterol-coupled ATPase activity *in vitro* [29]. The crystal structure herein provides a structural evidence of this peripheral sterol binding site where a sterol molecule makes direct contact with the conserved ABCG5_A540_ residue (**Fig. 2 & 5**). The sterol fits into an indented planar cleft within the upper region of the vestibule formed by the re-entry and the transmembrane helices. The captured cholesterol extends across the interfacial space, with its hydroxyl group forming a hydrogen bond with ABCG8_T430_, while the hydrocarbon tail remains straight and in contact with hydrophobic residues on ABCG5, such as L516, L537, I562, F565 and F567 (**Fig. 5D**). We also showed an inhibition of the cholesterol-binding activity by the A540F mutant (**Fig. 2C**). The K_D_ value is similar for both WT and mutant proteins, echoing the similar K_M_ for sterol-coupled ATPase activity in previous studies, given the mutant suppressed ATP hydrolysis by ∼90% [29]. Taken together with a 180° flipping of cholesterol by this mutant (**Fig. 5E**), current structural and functional data suggest a functionally defective conformation by this LOF mutant, inhibiting both sterol and ATP associations to the transporter. The Hill coefficient was estimated as ∼3, although no evidence so far shows whether the transporter bind one or multiple sterol molecules. Further structural and functional studies will be necessary to determine how these sterols bind ABCG5/G8.

In a separate experiment, cholesterol was docked to the TMD of previously published ABCG5/G8 structures (PDB ID: 5DO7 or 7JR7) [25, 33], where one of the top-3 predicted poses binds to the transporter in similar conformation as revealed by the crystal structure (**Fig. 3D/E**). The core steroid rings and the hydroxyl group matched in similar tilt, although the hydrocarbon tail varied slightly with minimal rotation. The latter may be explained by its relatively high mobility, such that the deviation was expected. Using a homology model containing the A540F mutation, the large aromatic group in position 540 introduced steric hindrance, consequently disabling cholesterol docking onto the observed position as shown in the crystal structure or predicted docking on WT protein (**Fig. 2 & 3**). In addition, cholesterol docking with the A540F mutant resulted in a loss of top-3 poses as seen on the WT protein. Altogether, past and present data support the existence of an orthogonal cholesterol binding site located around residue ABCG5_A540_, and the orthogonal sterol poses may be part of intermediate states of sterol-translocation trajectory along the TMD.

Via a protein sequence analysis of primarily mammalian ABCG transporters, we have identified a structural motif, “phenylalanine highway” along the TMH2 that consists of conserved aromatic residues (usually phenylalanine) (**Fig. 1**). This motif is flanked by two highly conserved aromatic residues (*i.e.*, positions F1 and F4), where position F4 appears to involve in sterol capturing from the cytoplasmic end of the membranes (**Suppl Fig. 1)** and the PH may provide a pathway for docked sterols to enter the dimer’s interface (**Fig. 3**). Although not fully defined, certain PH residues are believed to form a molecular clamp previously shown to mediate π-π interactions with sterol molecules [39,41,45,46]. As substrates for ABCG tend to be polycyclic with more or less an aromatic group that can interact with aromatic amino acids, the PH motif possibly serves as a conserved molecular clamp of ABCG sterol transporters (**Fig. 1C**). The clamp-like role of the lower PH terminus was recently suggested by structural studies of ABCG1 and ABCG2. The ATPase activity of ABCG1 was inhibited by ABCG1_F455A_ mutation [27], while ABCG2-driven drug efflux was downregulated by ABCG2_F439A_ mutation [46]. Interestingly, mutation ABCG2_F439Y_ or ABCG2_F439W_ did not hamper the efflux function, emphasizing the importance of aromatic residues in contributing to a substrate-binding route from the inner leaflet of the membranes [46]. In ABCG5/G8, while the clamp-like function of ABCG5_Y432_ has yet to be showcased, our docking analysis and a previous *in vivo* data suggest its biochemical nature as a key feature for the transporter function, presumably sterol binding. In this study, a double-mutant model (ABCG5_Y432F_/ABCG8_N564P_) revealed a loss of the ability to form a hydrogen bonding at the TMD dimer interface of the protein model (**Fig. 6B**) and a deeper cavity was formed to imitate ABCG1, yet the overall binding patterns were adversely affected (**Fig. 6D/E**). While the physiological impact of this double mutation has yet to be determined, such pair nonetheless created a blockade in the cavity of ABCG5/G8’s model. Since docking utilizes static protein models, widening of the cavity through mutation helped to mimic the effects of possible dynamic protein movement causing the subsequent fracture of the physical hindrance. This alteration in binding conformations and obstruction of the molecular clamp may help explain why *in vivo*, ABCG5_Y432A_/ABCG8_WT_ caused a significant decrease in biliary cholesterol efflux [25]. The mutations we had modelled retained the aromatic group; the *in vivo* data thus suggested a loss of the molecular clamp at the lower PH terminus. It is important to note that ABCG5_Y432_ is also part of the TMD polar relay network [23, 29]. Unlike the ABCG8 equivalent (ABCG8F461), ABCG5Y432 may play a dual role in the sterol binding/translocation and the allosteric regulation that couples ATPase and sterol-transport activities [29].

The upper PH terminus (*i.e.*, ABCG5Y424 and ABCG8F453) lies immediately under the hydrophobic valve (**Fig. 1D/E**), suggesting a cooperativity between PH’s F1 position and the hydrophobic valve in the sterol-transport function. ABCG5Y424, ABCG8F453 and ABCG1F447 all extend their side chains towards the valve’s aromatic residues, forming aromatic-aromatic interactions as suggested previously [27]. In ABCG1 and ABCG4, the di-phenylalanine valve may grant the possibility of aromatic interactions between the neighboring six phenylalanine residues. In contrast, the leucine and methionine from ABCG5/G8’s hydrophobic valve provide an asymmetric gating of sterol transport, as opposed to the symmetric valve in ABCG1 or ABCG4. In ABCG2, the lack of any phenylalanine within its di-leucine valve denies π-π interactions, although ABCG2F431 (PH’s F1) was shown to engage in drug-protein interactions with ABCG2 inhibitors [40]. Position F2 in the PH is moderately conserved **(Fig. 1A)**, although residues at this position has been proposed to involve sterol interactions, as suggested by decreased cholesterol efflux in the presence of ABCG1F448A mutant [32] or drug interaction in ABCG2 [47]. Interestingly, such a PH-like motif may not be limited to ABCG sterol transporters. We have observed similar phenylalanine spacing on the structures of other cholesterol-binding or -transport proteins, including ABCA1, ABCA7, SIDT1, SIDT2, PTCH1, and NPC1 (**Suppl Fig. 4**) [34,48– 51]. The PH structural motif may be more common than expected, and further studies will be needed to define its functional roles.

The docking experiments in this study have allowed us to analyze different binding profiles of cholesterol or stigmasterol on ABCG1 and ABCG5/G8 (**Fig. 3**). Previous *in vivo* studies showed that stigmasterol efflux by ABCG5/G8 was the fastest among dietary plant sterols [37]; hence stigmasterol was used to analyze phytosterol/plant sterol binding to ABCG transporters. Both proteins share cholesterol as a substrate yet deviate in phytosterol transporting abilities. ABCG5/G8 is a known phytosterol transporter, and each transporter-membrane interface has a distinct selectivity for one of the two docked sterols (**Fig. 3A/B**). In comparison, ABCG1 showed similar binding conformations between stigmasterol and cholesterol, with a decreased amount of stigmasterol binding poses on ABCG1 than ABCG5/G8. Together, our results indicate that sterol selectivity not only occurs on the opposite transporter-membrane interfaces but also differentiate ABCG5/G8 from other ABCG sterol transporters. In addition, our simulation showed the hydrocarbon tail of sterol molecules leading to the PH along with the TMD dimer interface, rather than the hydroxyl group as seen in previous cryo-EM structures of ABCG1 [27]. However, at least one docking pose on the ABCG8-dominant side horizontally points its hydroxyl group to the dimer interface, similar to that of a recent cryo-EM structure **(Suppl Fig. 5A)**. We speculate that the cholesterol orientation as seen in the cryo-EM structures may represent an intermediate cholesterol-binding state among the docked poses (**Fig. 5**).

Studies on cholesterol redistribution in lipid rafts have suggested a TMD sterol access site in ABCG1 (**Fig. 4**) [16,28,52]. Theoretically, ABCG1 intakes sterols from enriched lipid microdomains, relocating them towards the surrounding sterol-deprived membranes. The docking pattern follows the amphipathic nature of the lipid bilayers with cholesterol’s polar head localizing around the cytoplasmic matrix **(Fig. 3)**, where. cholesterol’s hydroxyl group is believed to generate hydrogen bonds with the surrounding polar lipid regions, orienting the sterols head towards the hydrophilic region [53]. Our docking analysis suggests ABCG1’s apparent preference for the inner leaflet sterols [16], where the transporters then translocate cholesterol to the upper leaflet, similar to the recently proposed mechanism [26, 32]. In ABCG5/G8, both crystallographic and docking results indicated that sterol binding is not limited to the TMD cavity open towards to the cytoplasmic side, but an orthogonal sterol-binding state is a possible intermediate state to drive sterol efflux. As the TMD dimer interface in ABCG5/G8 is much narrower than that in ABCG1, it is possible that ABCG5/G8 similarly uptakes sterol substrates as ABCG1. Still, sterols would need to undergo a series of yet-to-be characterized movements from their longitudinal orientations to an orthogonal binding state as shown in the cholesterol-bound structure (**Fig. 2 &3**). Among ABCG5/G8 substrates, plant sterols may show higher binding preferences as suggested by stigmasterol poses (**Fig. 3**).

In conclusion, our crystallographic and computational analysis have highlighted possible regions of sterol binding and interactions on ABCG5/G8. This study shows a conserved phenylalanine highway motif on TMH2 that may play an essential role in substrate recruitment and binding and a crystal structure revealing an orthogonal cholesterol bound on the ABCG5-dominant transporter-membrane interface, consistent with the leading cholesterol poses predicted by molecular docking. The docking analysis of cholesterol and stigmasterol also suggested the asymmetric sterol-binding potential on ABCG5- and ABCG8-dominant transporter-membrane interfaces, and difference in sterol-docking patterns may contribute to different substrate specificity between ABCG1 and ABCG5/G8. Further structural and functional studies will reveal more insight into sterol selectivity and translocation trajectory by ABCG sterol transporters.

## 4. MATERIALS AND METHODS

### 4.1 Protein purification and crystallization

For purification of human WT ABCG5/G8 from *Pichia pastoris* yeast, we have followed the procedures based on our previously reported protocol [25]. With minor modifications, the procedures are briefly described in the following. For each purification, six litres of yeast cell culture were used for ∼200 mL of the microsomal membrane preparation at the total protein content of ∼10 mg/mL. The membranes were solubilized by equal volume of the solubilization buffer (50 mM Tris-HCl, pH 8.0, 100 mM NaCl, and 10% glycerol, 2% (w/v) β-dodecyl maltoside (β-DDM, Anatrace), 0.5% (w/v) cholate (Sigma-Aldrich), 0.25% (w/v) cholesteryl hemisuccinate Tris (CHS-Tris, Anatrace), 5 mM imidazole, 5 mM β-mercaptoethanol (β-ME), 2 μg/ml leupeptin, 2 μg/ml pepstatin A, 2 mM PMSF). Insoluble membranes were removed by ultracentrifugation, and the solubilized supernatant was treated with a final concentration and 0.1 mM Tris (2-carboxylethylcarboxyethyl) phosphine (TCEP). Following that, tandem affinity chromatography was performed. The soluble membrane proteins were separated using a 20-mL nickel-nitrilotriacetic acid (Ni-NTA) column. Then, the peak fractions of ABCG5/G8 from the Ni-NTA eluates were mixed with the equal amount of CBP column wash buffer (50 mM HEPES, pH 7.5, 100 mM NaCl, 0.1% (w/v) β-DDM, 0.05% (w/v) cholate, 0.01% (w/v) CHS (Steraloids), 0.1 mM TCEP ,1 mM CaCl_2_, 1 mM MgCl_2_) and then loaded onto a 5-mL CBP column. The CBP column was washed sequentially to exchange detergents to DMNG, and ABCG5/G8 proteins were finally collected using an elution buffer (50 mM HEPES, pH 7.5, 300 mM NaCl, 2 mM EGTA, 0.1% (w/v) DMNG, 0.05% (w/v) cholate, 0.01% (w/v) CHS, 1 mM TCEP). The N-linked glycans and the CBP tag were cleaved by endoglycosidase H (Endo H, ∼0.2 mg per 10–15 mg purified protein, New England Biolabs) and HRV-3C protease (∼2 mg per 10–15 mg purified proteins) for overnight at 4 °C. The CBP tag-free proteins were further purified by gel filtration chromatography using an ÄKTA Purifier and a Superdex 200 30/100 GL column in the buffer containing 10 mM HEPES, pH 7.5, 100 mM NaCl, 0.1% (w/v) DMNG, 0.05% (w/v) cholate, 0.01% (w/v) CHS. The peak fractions were pooled together and treated by reductive methylation. The methylated proteins were relipidated by overnight incubation with 0.5 mg/ml DOPC: DOPE (3:1, w/w) and further purified by a 2-mL Ni-NTA column, followed by iodoacetamide treatment for cysteine alkylation. The relipidated proteins were treated with 1 mM AMPPNP (sodium salt, Roche) for overnight at 4 °C and desalted with a PD-10 column. The desalted and lipidated proteins were incubated overnight with cholesterol (prepared in isopropanol) to a final concentration of ∼20 μM, and protein precipitants were removed by ultracentrifugation for 10 min at 4 °C. Using 100 kDa cutoff filters, the supernatants were concentrated to a final protein concentration of 30-50 mg/ml and crystallized within a week.

Protein crystal growth was carried out by bicelle crystallization as described previously. Briefly, the cholesterol-treated ABCG5/G8 proteins were reconstituted into 10% DMPC/CHAPSO bicelles (lipids and detergents in a 3:1 (w/w) ratio, with the lipids containing 5 mol% cholesterol and 95 mol% DMPC). After that, the proteins and bicelle stock were gently mixed in a 1:4 (v/v) ratio, resulting in a final protein concentration of ∼10 mg/ml. At 20 °C, the crystallization was done in a hanging-drop vapor diffusion technique by mixing protein/bicelle preparation with equal-volume crystallization reservoir solution containing 1.8–2.0 M ammonium sulfate, 100 mM MES pH 6.5, 2–5% PEG400, and 1 mM TCEP. Protein crystals were harvested within 1-2 weeks of incubation by first submerging the crystals with 0.2 M sodium malonate and then flash-freezing the crystals in liquid nitrogen with Mitegen cryoloops.

### 4.2 Data collection of X-ray diffraction, structural determination and refinement

X-ray diffraction data were collected using beamlines 19-ID on a Quantum 315r detector at the Advanced Photon Source (APS). Scaling and integration of three datasets were carried out by HKL2000 software package, and the final data was scaled between 30Å and 4Å resolutions to ensure I/σ >1 [54]. Using Phaser program, we obtain the phase information and a structural solution by molecular replacement using a cryo-EM structure (PDB ID: 7JR7) as the initial template [33, 55]. Before model refinement by Phenix, the Phaser solution was corrected at the Walker A, the Walker B, and the Signature motifs by the more defined registry in the crystal structure (PDB ID: 5DO7) [25, 56]. The initial refinement resulted in a refined model with Rfree and Rwork of 0.338 and 0.309, respectively. Close inspection of the Fo-Fc map, two orthogonal electron densities that clearly include the nature of polycyclic rings were observed between the crystallographic dimer interface and on the ABCG5-dominant side where each electron density is within the van der Waals’ distance from ABCG5_A540_. No such electron densities were observed on ABCG8-dominant side. We thus modeled a cholesterol molecule per electron density, and we refined the structure in the presence of one cholesterol molecule on each ABCG5/G8 heterodimer by testing a series of sterol orientations using the program COOT [57]. Following our final refinement, Rfree and Rwork were determined to be 0.302 and 0.244, respectively. The model quality was validated by MolProbity as implemented in Phenix [56, 58]. As a note, given the proteins were co-crystallized with AMPPNP, we were not able to observe obvious electron densities for the nucleotides from the current datasets at the current resolution.

### 4.3 Protein sequence analysis

All FASTA files were acquired using PSI-BLAST (position-specific iterated basic local alignment search tool) as implemented in the NCBI protein sequence database. All queries were acquired such that the organism belonged to the Mammalia family, and all partial and incomplete queries were filtered out. FASTA sequences were then surveyed one by one to ensure that the search criteria were executed properly, and no incomplete sequences were passed. FASTA sequences were acquired for ABCG1, ABCG2, ABCG4, ABCG5, and ABCG8 (P45844, Q9UNQ0, Q9H172, Q9H222, Q9H221) with the following PSI-BLAST search query: *"ABCXY"[Gene] AND "Mammalia"[Organism] NOT "partial"[All Fields],* Whereby X denotes the ABC subfamily, and Y denotes the member protein. Including 250-300 species, all alignments and analyses were run locally using the Bio Python repository [59]. A multiple sequence alignment (MSA) was conducted using all the FASTA sequences using a BLOSUM62 substituation matrix [60], ClustalW [61] or PROMALS3D [62]. To score conservation, alignment files were made of the FASTA sequence library using the Multiple Sequence Comparison by Log-Expectation (MUSCLE) tool [63]. To generate a consensus sequence, a “dumb consensus” was conducted with a default threshold of 70% and a required multiple of 1. Consensus sequences were acquired for mammalian ABCG1, ABCG2, ABCG4, ABCG5, and ABCG8. Consensus sequences for each protein subfamily and member were compared to one another to determine highly conserved amino acids and amino acid sequences. For example, the amino acids present in the consensus sequence of ABCG1 was individually aligned to ABCG2, ABCG4, ABCG5, and ABCG8. Conservation scores above the set threshold (80%) were extracted for analytical purposes. An 80% similarity threshold, as opposed to 70%, was used using AMAS method of multiple sequence alignment because this analysis compared two consensus sequences together, 80% was chosen arbitrarily to ensure the amino acids chosen were highly similar without extracting ambiguous amino acid [64]. For further analysis, the results from the conservation scores were compared to each other (*i.e.,* the conservation scores of ABCG5 and ABCG2 was compared to conservation scores of ABCG5 and ABCG8). This was done across all comparisons to generate a comprehensive list of the amino acids that are conserved across all subfamily and member combinations. The same procedure was used to analyze ABCA1, ABCA7, SIDT1, SIDT2, PTCH1, and NPC1, of which experimental structures have been reported [48–51].

### 4.4 Homology Modelling

UCSF Modeller was used for homology modelling [65]. The sequence alignment was created between the template and the mutant sequences using PROMALS3D [62]. The template is the currently accepted PDB ID: 5DO7 in conjunction with X-ray data to create a more accurate model with a better energy DOPE (discrete optimized protein energy) score. The mutant sequences included ABCG5_Y432F_, ABCG8_N564P_ and ABCG5_A540F_. After the alignment, UCSF Modeller was used to generate 400 models and to rank them based on their DOPE scores. The best model was taken and visualized using PyMOL or UCSF Chimera. For the ATP-bound model, the cryo-EM structure of ATP-bound ABCG2 E211Q (PDB ID: 6HBU) was used as the template [41]. 100 models were generated with MODELLER, and the model with the best DOPE score was used for the sterol docking experiments in this study. All models were evaluated using MolProbity web server [58] to ensure no severe geometric violation in the Ramachandran plot [66].

### 4.5 Protein and ligand model preparation for docking

Protein models ABCG1, ABCG2, ABCG5/G8, PTCH1, NPC1, ABCA1 and PDR5 gathered from RCSB PDB database, PDB ID: 7OZ1/7R8D, 6VXI/6VXF/6VXJ, 5DO7/7R8E/7JR7, 6OEU, 6UOX, 5XJY, 7P03, respectively. ABCG4, White protein, ABCA7, SIDT1/2 structures were downloaded from AlphaFold2 predictions [25–27,33,34,39,48–51]. Regions of interest in these proteins had very high confidence scoring (pLDDT>90). Visualization of the structures done through PyMOL and Chimera. Double mutant ABCG5_Y432F_ and ABCG8_N564P_ and ABCG5_A540F_ were induced and modelled in MODELLER based on PDB ID: 5DO7. The proteins docked (ABCG1, ABCG5/G8, ATP-bound, ABCG5/G8 mutants) were cut at the elbows, excluding the nucleotide-binding domain due to the large size of the full protein (>999 amino acids). Cholesterol was gathered from PDB ID: 7R8D, while stigmasterol was obtained in PubChem [27]. ABCG2 ligands used for docking (imatinib and SN38) were previously modelled within the structure [39, 40]. Dunbrack 2010 rotamer library [67] and ANTECHAMBER [68] were used in dockprep charge and hydrogen bond additions.

### 4.6 Molecular Docking

UCSF Dock6.9 [38] was utilized through SBGrid [69] to perform molecular docking. UCSF Chimera [70] was used as the graphic software for the simulated models. Protein binding site was determined by “sphgen” estimation specified at 10 Å distance from the ligand, located roughly in the centre of the protein in ABCG transporters. The “gridbox” and “grid” were generated and analyzed prior to the dock. To avoid potential mixing-up, we first performed docking with each sterol ligand on either side of transporter-membrane interface, whose results were consistent with those when placing the ligands at the centre of the TMD interface. Grid was built at a distance of 8 Å from the selected “sphgen” spheres, and ligand’s energy was minimized through minor conformational changes. A flexible docking procedure was then followed by calculation of 1000 or 5000 orientations, yielding no difference in results. The best scoring poses were compiled and individually viewed through UCSF Chimera’s viewdock tool, these were then ranked by van der Waals energy (vdw_energy) and root mean square deviation (RMSD). Interacting amino acids were identified and plotted by both Chimera and PyMOL [70].

To validate our protocol for sterol docking on ABCG1 or ABCG5/G8, we performed a control experiment using ABCG2 and three of its known drugs (i.e., imatinib, and SN38) [39]. Using a previously published ABCG2 structure (PDB ID: 6FFC, 6VXH and 6VXJ), ABCG2 was docked with the drug ligand (one at a time) and weighed again against the originally modelled pose [40]. The simulated results showed consistent orientation of the drugs between predicted and experimental data, and the best pose for each drug fits the originally modelled ligand with the best RMSD and the lowest van vdw_energy value. The “best pose” in this study was therefore defined as the lowest van der Waal’s (vdw) energy, which was used to rank all of the docking results.

### 4.7 Cholesterol binding assay

A direct cholesterol-binding assay using detergent-purified ABCG5/G8 was developed based on the previously described [71]. Briefly, 0.24µM of His-tagged WT ABCG5/G8, the mutant or the control protein (*i.e.*, BSA) was incubated with 0-4 µM of cholesterol that was prepared in the presence of [^3^H]Cholesterol with a specificity of 110cpm/fmol (Perkein Elmer) in 50 µl of binding buffer containing 50mM Tris-HCl pH7.5, 100mM NaCl, and 0.05%(w/v) β-DDM (Anatrace), 0.05% (w/v) sodium cholate (Sigma-Aldrich), 1mM TCEP, 30µl of pre-equilibrated Ni-NTA bead (QIAGEN) on ice for overnight. The supernatant was removed by centrifugation at 600 rpm for 1min in a bench-top centrifuge. The bead was extensively washed with 600 µl of ice-cold binding buffer plus 0.01% (w/v) CHS (Steraloids) for 4 times. To elute protein, 600 µl of binding buffer plus 0.01% (w/v) CHS and 200mM imidazole was added and incubated with the washed bead on ice for 30min. The bound [^3^H]Cholesterol was quantified by measuring the radioactivity of 500µl of elution fraction in a liquid scintillation counter (Perkin Elmer Tri-Carb 4910TR).

## 5. Data Availability

Diffraction data have been deposited in PDB under the accession code 8CUB.

## AUTHOR CONTRIBUTIONS

DF carried out docking analysis of ABCG transporters and other cholesterol-binding proteins; JYL performed crystallization and collected X-ray diffraction data; FR processed the X-ray data and determined the crystal structure. DF, MR and AS performed sequence analysis; MR and LD generated homology models. YY, JFC and JYL designed the cholesterol-binding assay; YY optimized the assay and analyzed the data. JYL oversaw the project, and DF, MR, FR, SS and JYL wrote the manuscript. All authors have read and agreed to the submitted version of the manuscript.

## ACKNOWLEDGEMENTS

We thank the technical help from the staff of both the Common Equipment & Technical Services and the Protein Biophysics Core Facilities, as well as the critical feedback from Drs. Thomas Stockner, Gregory Graf, and Ms. Annahat Kochhar. We also thank Aiman Zein, Angelica Venes and Toka Hussein for technical assistance. This work is partially based on the theses that were submitted to fulfill in part the requirement for the degrees of Honors Bachelor of Science (DF & MR). FR is partially supported by an Academia Sinica-University of Ottawa Travel Support for Research Projects. The Advanced Photon Source is a US Department of Energy Office of Science User Facility operated for the Department of Energy Office of Science by Argonne National Laboratory (DE-AC02-06CH11357).

## ABREVIATIONS

ABC: Adenosine-triphosphate-binding cassette
NBD: Nucleotide binding domain
TMD: Transmembrane domain
TMH: Transmembrane helix
RCT: Reverse cholesterol transport
PH: Phenylalanine Highway

---

**Supplemental Figure 1.**
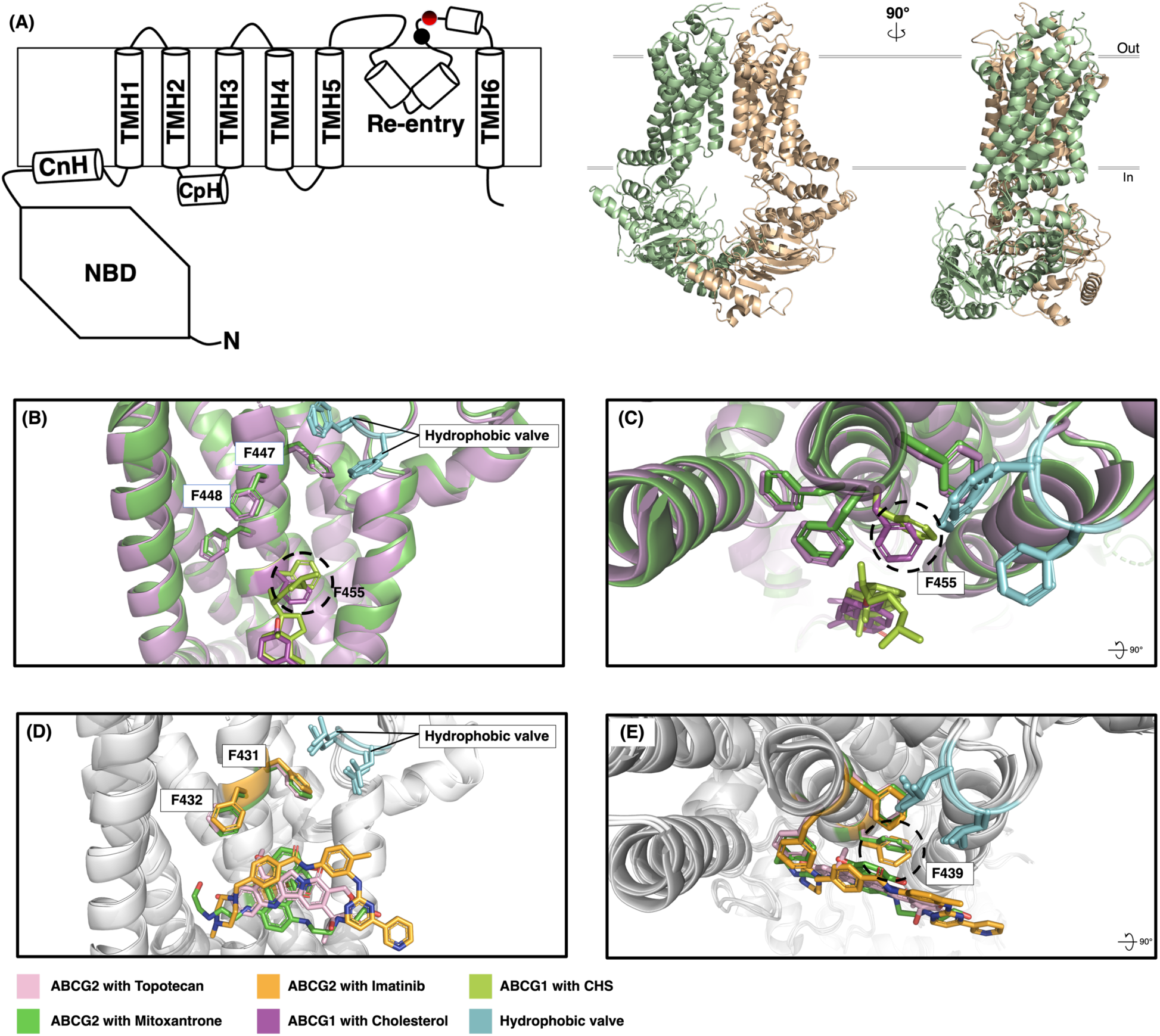
Topological representation of the ABCG5/G8 and Superposition of the phenylalanine highway on ABCG1 and ABCG2. **(A)** *Left,* Topological representation of the ABCG5/G8 crystal structure (PDB ID: 8CUB) is depicted in empty cylinders, showing the transmembrane structural segments of ABCG5/G8 including connecting helixes (CnH), TM helixes, coupling helices (CpH), and re-entry helixes. *Right,* The cartoon illustration of the ABCG5/G8 heterodimer with ABCG5 in pale-green and ABCG8 in tan colors. The N-glycosylation sites are shown in solid circles: ABCG5_N592_ or ABCG8_N619_ in red and black, ABCG5_N584_ in black. **(B)** Side view of ABCG1 (PDB ID: 7OZ1), showing the difference in height of CHS (green) and cholesterol (purple). **(C)** Top-down view ABCG1 bound to CHS (PDB ID: 7OZ1, green) aligned with ABCG1 bound to cholesterol (7R8D, purple) emphasizing the difference in F455 rotation when interacting with different length ligands. Rotation occurs when longer ligands (such as CHS) reach deep enough into the transmembrane cavity, causing the alpha carbon and phenyl side chain of F455 to rotate. F447 is also seen stacked under hydrophobic valve. **(D)** Side view of ABCG2, showing the difference in localization of Imatinib (orange), Topotecan (pink), Mitoxantrone (green). **(E)**Top-down view of ABCG2 bound with Imatinib (PDB ID: 6VXH, orange), Topotecan (PDB ID :6VXJ, pink) and Mitoxantrone (PDB ID: 6VXI, green). Rotation of F439 is seen when substrates are deep enough into the cavity to interact with the residue. Contrarily, when they are not, such rotation events do not happen, as observed with Imatinib.

**Supplemental Figure 2.**
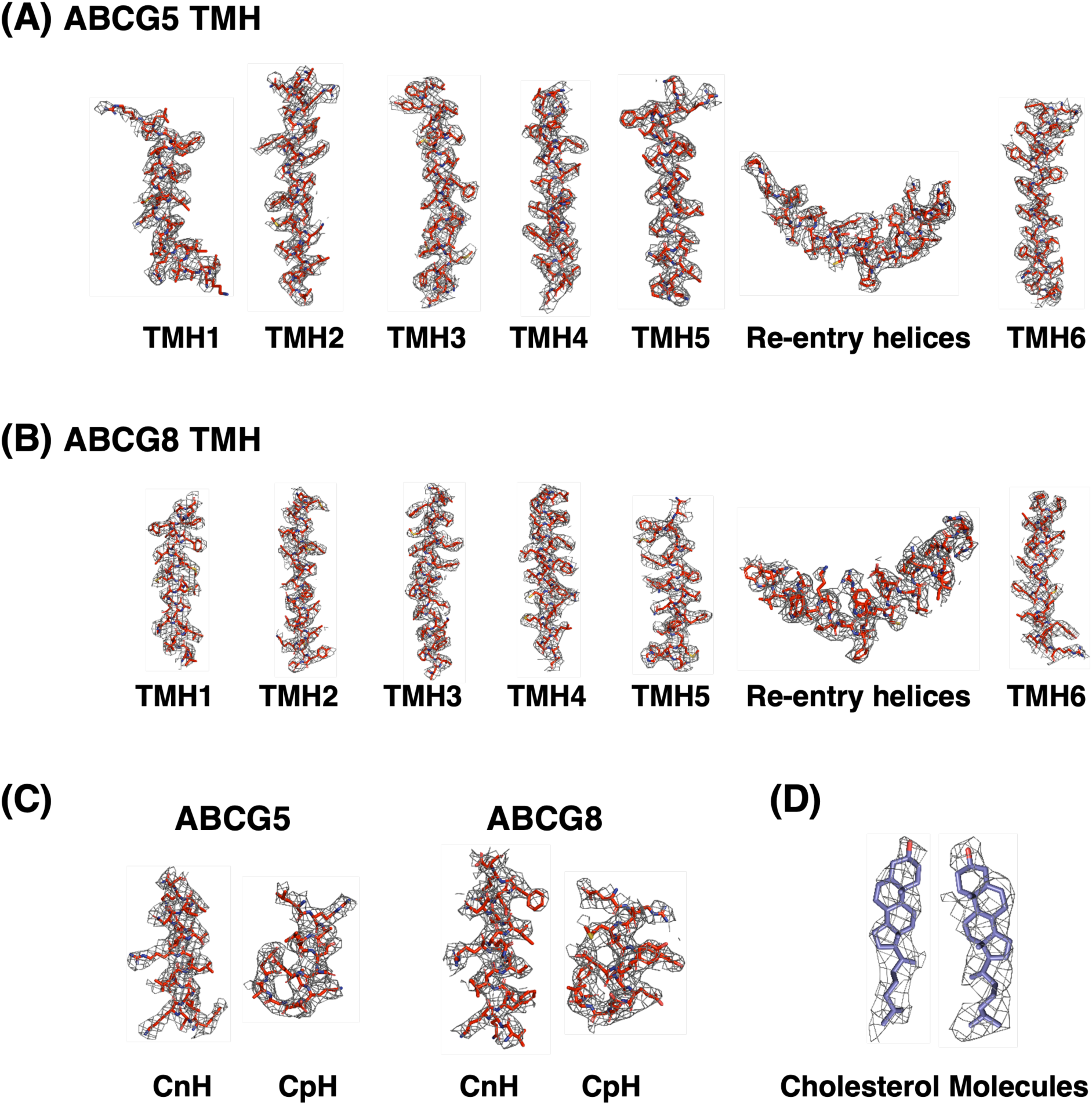
The fitting of the structural model in the electron densities (2Fo-Fc omit map). The representative segments of the human ABCG5/G8 proteins, including transmembrane (A/B), connecting (C), coupling helixes (C), and cholesterol molecules (D) are shown separately for G5 and G8 chains in the electron density with a mesh contoured at 1 σ and the thickness of 1.5 as implemented in PyMOL.

**Supplemental Figure 3.**
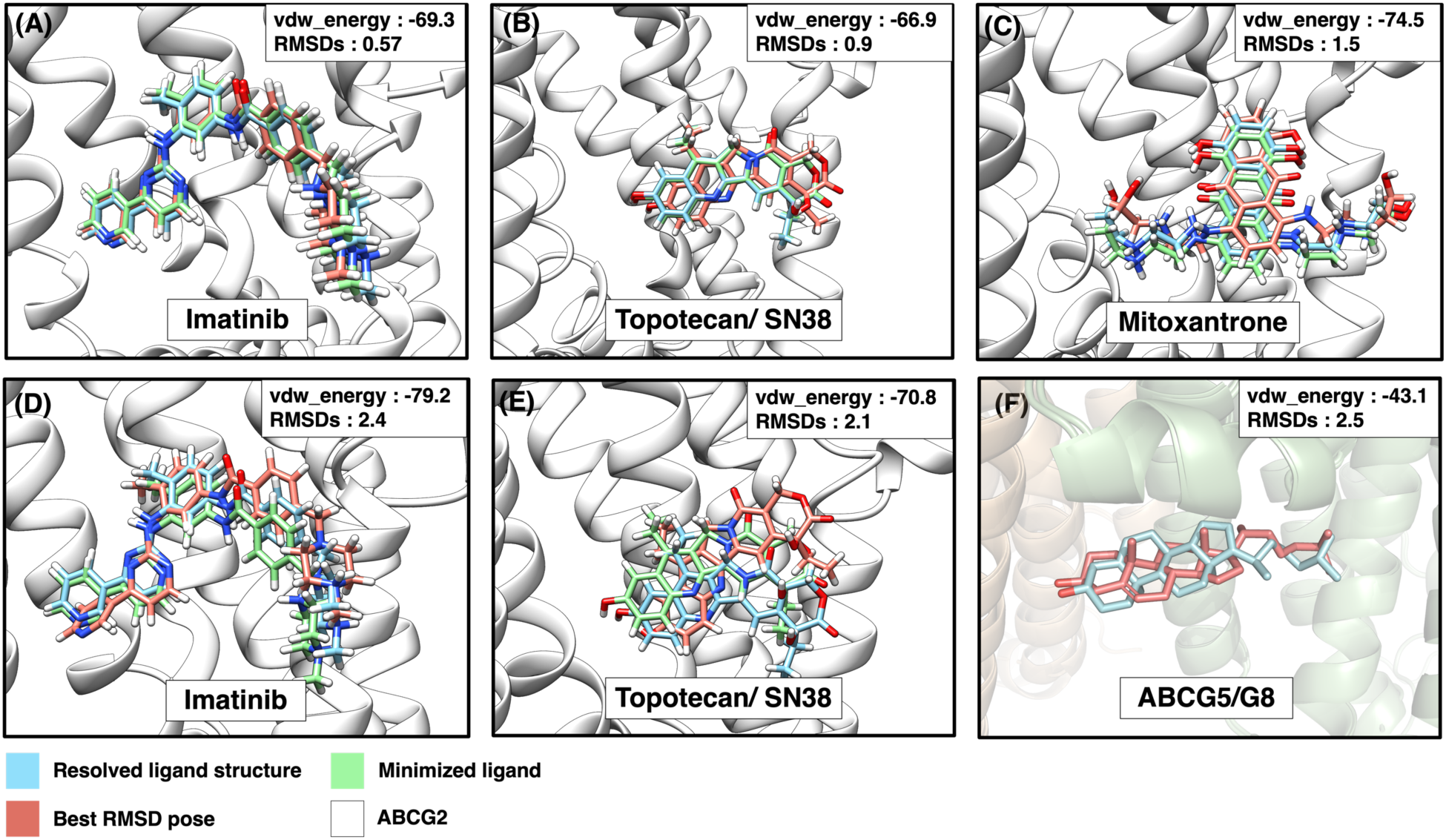
Resolved ligand structures compared to dock prediction in ABCG2 (A-E) and ABCG5/G8 (F). (**A-C**) Redocking of resolved ligands on ABCG2 (blue): Topotecan/SN38 (PDB ID :6VXJ), Mitoxantrone (PDB ID: 6VXI), and Imatinib (PDB ID: 6VXH). Green highlights the minimized rigid conformation as calculated by Dock6.9. Orange highlights the best predicted pose. Blue ligands were observed in the experimentally determined structures. RMSD of all superpositions is (0.57Å – 1.5Å). (**D/E**) Ligands were aligned and docked to ABCG2 (PDB ID: 6FFC). Pink highlights the best RMSD predicted pose. The RMSD of all superpositions is (2.1Å – 2.4Å). (**F**) Redocking of cholesterol ligand on ABCG5/G8 (RMSD 2.5Å, present study). See details in Materials and Methods.

**Supplemental Figure 4.**
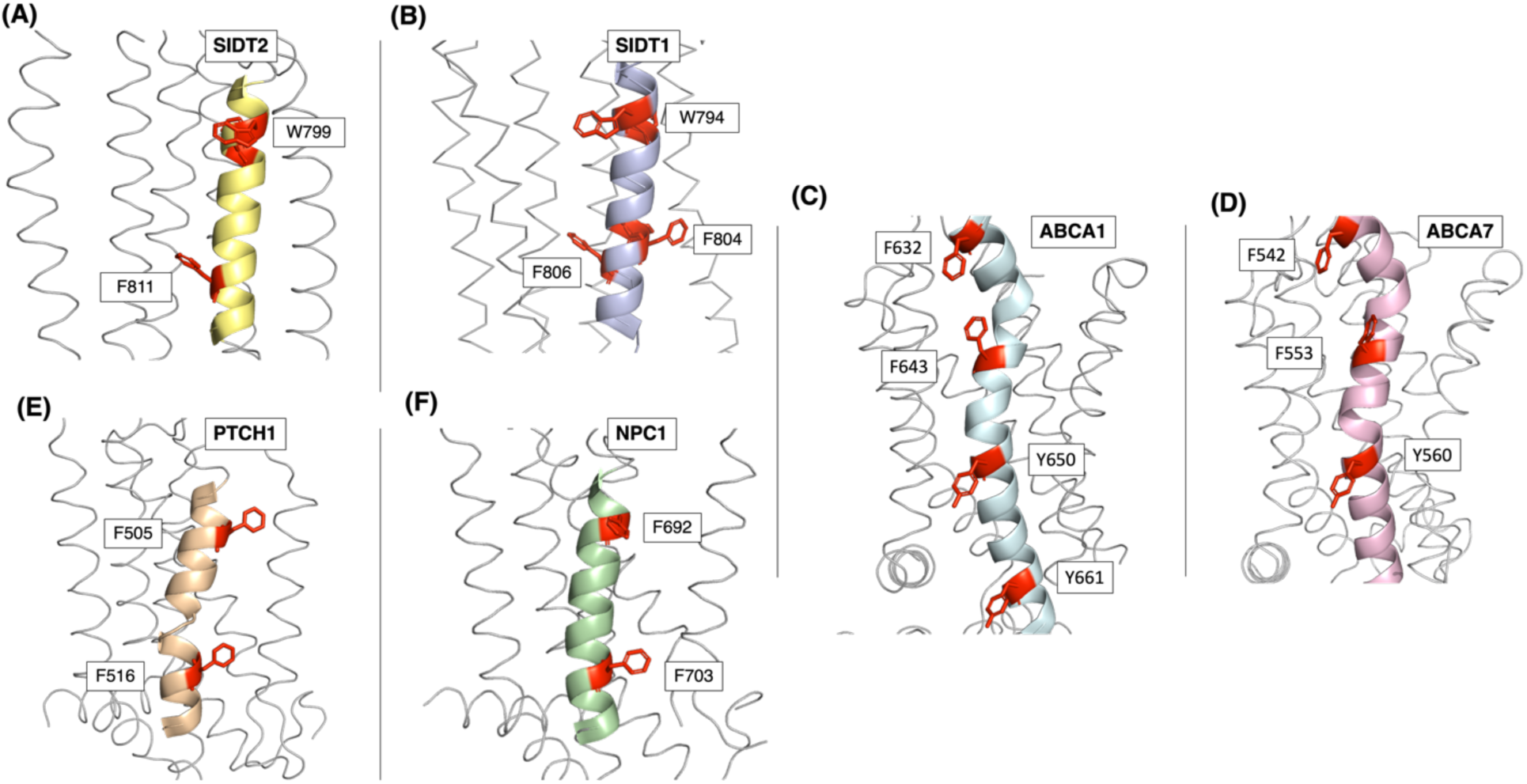
Aromatic amino acid pattern in non-ABCG lipid transporters. Conserved aromatic residues along the putative (or observed) sterol-binding/interacting TMH (cartoon presentation) show a similar pattern of phenylalanine highway as observed in ABCG transporters. SIDT2 (**A**, yellow), SIDT1 (**B**, dark blue), ABCA1 (**C**, light blue), ABCA7 (**D**, pink), PTCH1 (**E**, orange) and NPC1 (**F**, green). The aromatic amino acids: phenylalanine, tyrosine and tryptophan are highlighted in red sticks. Only the most central and luminal helices are modelled in detail.

**Supplemental Figure 5.**
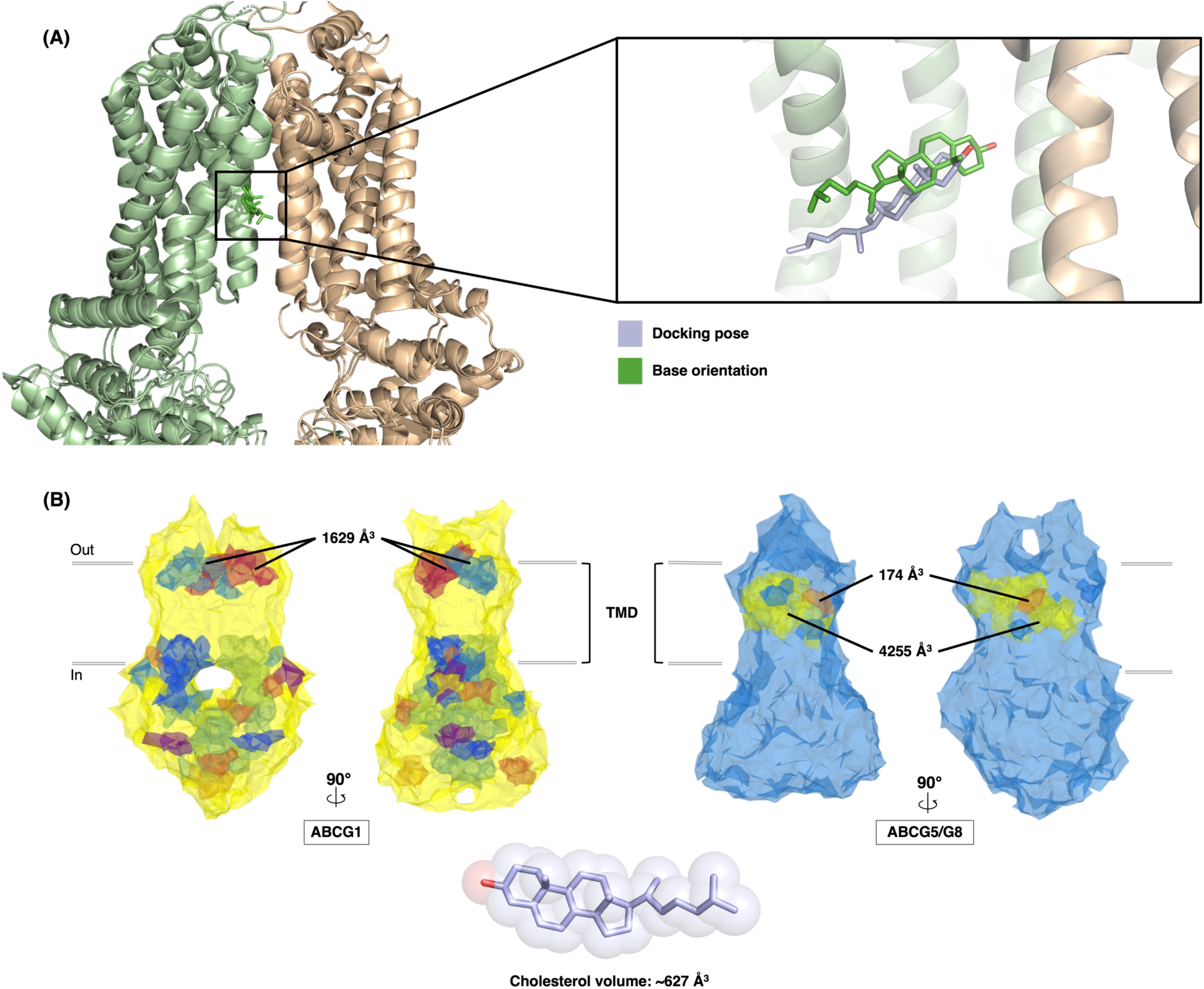
Cavities and binding regions in ABCG structures. **(A)** *Left.* ABCG5/G8 (PDB ID: 5DO7) aligned with cryo-EM structure (PDB ID: 7R8D). *Right.* Comparison of cholesterol docking pose (light blue) with the cholesterol (green) in the cryo-EM structure [28]. **(B)** Volumes of transmembrane domain (TMD) cavities found in the of ATP-bound ABCG1 (yellow, PDB ID: 7R8E) and ABCG5/G8 (blue) calculated by MOLE2.5. *Left*, two cavities in ABCG1 found near the interface between the TMD and extracellular domain (ECD). These cavities (1629Å^3^ each) curve over the central canal of the protein. *Right*, one large cavity found spanning throughout the TMD. This cavity represents a conglomeration of sphere-shaped cavities connected by thin bridging regions. Relevant cavities in the central canal of the protein are represented as spherical and estimated to be near the size of ∼200Å^3^. *Bottom*, the volume of a cholesterol molecule is 627Å^3^.

## REFERENCES

[1] W. Nan, L. Debin, C. Wengen, M. Fumihiko, T.A.R., ATP-binding cassette transporters G1 and G4 mediate cellular cholesterol efflux to high-density lipoproteins, Proc. Natl. Acad. Sci. 101 (2004) 9774–9779. https://doi.org/10.1073/pnas.0403506101.

[2] A.M. Vaughan, J.F. Oram, ABCG1 Redistributes Cell Cholesterol to Domains Removable by High Density Lipoprotein but Not by Lipid-depleted Apolipoproteins *, J. Biol. Chem. 280 (2005) 30150–30157. https://doi.org/10.1074/jbc.M505368200.

[3] L. Yu, S. Gupta, F. Xu, A.D.B. Liverman, A. Moschetta, D.J. Mangelsdorf, J.J. Repa, H.H. Hobbs, J.C. Cohen, Expression of ABCG5 and ABCG8 Is Required for Regulation of Biliary Cholesterol Secretion *, J. Biol. Chem. 280 (2005) 8742–8747. https://doi.org/10.1074/jbc.M411080200.

[4] D.L. Austin, Y. Weidong, A.L. V., K. Tammy, G. Yongming, R.A. K., R.D. D., A multidrug resistance transporter from human MCF-7 breast cancer cells, Proc. Natl. Acad. Sci. 95 (1998) 15665–15670. https://doi.org/10.1073/pnas.95.26.15665.

[5] J.D. Allen, A. van Loevezijn, J.M. Lakhai, M. van der Valk, O. van Tellingen, G. Reid, J.H.M. Schellens, G.-J. Koomen, A.H. Schinkel, Potent and Specific Inhibition of the Breast Cancer Resistance Protein Multidrug Transporter in Vitro and in Mouse Intestine by a Novel Analogue of Fumitremorgin C1, Mol. Cancer Ther. 1 (2002) 417–425.

[6] Y. Namba, C. Sogawa, Y. Okusha, H. Kawai, M. Itagaki, K. Ono, J. Murakami, E. Aoyama, K. Ohyama, J. Asaumi, M. Takigawa, K. Okamoto, S.K. Calderwood, K. Kozaki, T. Eguchi, Depletion of Lipid Efflux Pump ABCG1 Triggers the Intracellular Accumulation of Extracellular Vesicles and Reduces Aggregation and Tumorigenesis of Metastatic Cancer Cells , Front. Oncol. . 8 (2018). https://www.frontiersin.org/article/10.3389/fonc.2018.00376.

[7] C. Tian, D. Huang, Y. Yu, J. Zhang, Q. Fang, C. Xie, ABCG1 as a potential oncogene in lung cancer, Exp Ther Med. 13 (2017) 3189–3194. https://doi.org/10.3892/etm.2017.4393.

[8] M.-H. Lee, K. Lu, S. Hazard, H. Yu, S. Shulenin, H. Hidaka, H. Kojima, R. Allikmets, N. Sakuma, R. Pegoraro, A.K. Srivastava, G. Salen, M. Dean, S.B. Patel, Identification of a gene, ABCG5, important in the regulation of dietary cholesterol absorption, Nat. Genet. 27 (2001) 79–83. https://doi.org/10.1038/83799.

[9] K.E. Berge, T. Hui, G.G. A., Y. Liqing, G.N. V., S. Joshua, K. Peter, S. Bei, B. Robert, H.H. H., Accumulation of Dietary Cholesterol in Sitosterolemia Caused by Mutations in Adjacent ABC Transporters, Science (80-. ). 290 (2000) 1771–1775. https://doi.org/10.1126/science.290.5497.1771.

[10] I. Meurs, B. Lammers, Y. Zhao, R. Out, R.B. Hildebrand, M. Hoekstra, T.J.C. Van Berkel, M. Van Eck, The effect of ABCG1 deficiency on atherosclerotic lesion development in LDL receptor knockout mice depends on the stage of atherogenesis, Atherosclerosis. 221 (2012) 41–47. https://doi.org/https://doi.org/10.1016/j.atherosclerosis.2011.11.024.

[11] T.M. Do, M.-S. Noel-Hudson, S. Ribes, C. Besengez, M. Smirnova, S. Cisternino, M. Buyse, F. Calon, G. Chimini, H. Chacun, J.-M. Scherrmann, R. Farinotti, F. Bourasset, ABCG2- and ABCG4-Mediated Efflux of Amyloid-β Peptide 1-40 at the Mouse Blood-Brain Barrier, J. Alzheimer’s Dis. 30 (2012) 155–166. https://doi.org/10.3233/JAD-2012-112189.

[12] A.A. Zein, R. Kaur, T.O.K. Hussein, G.A. Graf, J.-Y. Lee, ABCG5/G8: a structural view to pathophysiology of the hepatobiliary cholesterol secretion, Biochem. Soc. Trans. 47 (2019) 1259–1268. https://doi.org/10.1042/BST20190130.

[13] Y. Liqing, H.R. E., L.-H. Jia, von B. Klaus, L. Dieter, C.J. C., H.H. H., Disruption of Abcg5 and Abcg8 in mice reveals their crucial role in biliary cholesterol secretion, Proc. Natl. Acad. Sci. 99 (2002) 16237–16242. https://doi.org/10.1073/pnas.252582399.

[14] S.B. Patel, A. Honda, G. Salen, Sitosterolemia: exclusion of genes involved in reduced cholesterol biosynthesis, J. Lipid Res. 39 (1998) 1055–1061. https://doi.org/10.1016/S0022-2275(20)33874-8.

[15] E. Pandzic, I.C. Gelissen, R. Whan, P.J. Barter, D. Sviridov, K. Gaus, K.-A. Rye, B.J. Cochran, The ATP binding cassette transporter, ABCG1, localizes to cortical actin filaments, Sci. Rep. 7 (2017) 42025. https://doi.org/10.1038/srep42025.

[16] O. Sano, S. Ito, R. Kato, Y. Shimizu, A. Kobayashi, Y. Kimura, N. Kioka, K. Hanada, K. Ueda, M. Matsuo, ABCA1, ABCG1, and ABCG4 are distributed to distinct membrane meso-domains and disturb detergent-resistant domains on the plasma membrane, PLoS One. 9 (2014). https://doi.org/10.1371/journal.pone.0109886.

[17] S. Savary, F. Denizot, M.-F. Luciani, M.-G. Mattei, G. Chimini, Molecular cloning of a mammalian ABC transporter homologous to Drosophila white gene, Mamm. Genome. 7 (1996) 673–676. https://doi.org/10.1007/s003359900203.

[18] D.D. Bojanic, P.T. Tarr, G.D. Gale, D.J. Smith, D. Bok, B. Chen, S. Nusinowitz, A. Lövgren-Sandblom, I. Björkhem, P.A. Edwards, Differential expression and function of ABCG1 and ABCG4 during development and aging, J. Lipid Res. 51 (2010) 169–181. https://doi.org/10.1194/M900250-JLR200.

[19] T.Q. de Aguiar Vallim, E. Lee, D.J. Merriott, C.N. Goulbourne, J. Cheng, A. Cheng, A. Gonen, R.M. Allen, E.N.D. Palladino, D.A. Ford, T. Wang, Á. Baldán, E.J. Tarling, ABCG1 regulates pulmonary surfactant metabolism in mice and men[S], J. Lipid Res. 58 (2017) 941–954. https://doi.org/https://doi.org/10.1194/jlr.M075101.

[20] D. Sag, C. Cekic, R. Wu, J. Linden, C.C. Hedrick, The cholesterol transporter ABCG1 links cholesterol homeostasis and tumour immunity, Nat. Commun. 6 (2015) 6354. https://doi.org/10.1038/ncomms7354.

[21] G.A. Graf, L. Yu, W.-P. Li, R. Gerard, P.L. Tuma, J.C. Cohen, H.H. Hobbs, ABCG5 and ABCG8 Are Obligate Heterodimers for Protein Trafficking and Biliary Cholesterol Excretion*, J. Biol. Chem. 278 (2003) 48275–48282. https://doi.org/https://doi.org/10.1074/jbc.M310223200.

[22] C. Yang, L. Yu, W. Li, F. Xu, J.C. Cohen, H.H. Hobbs, Disruption of cholesterol homeostasis by plant sterols, J. Clin. Invest. 114 (2004) 813–822. https://doi.org/10.1172/JCI22186.

[23] N. Khunweeraphong, J. Mitchell-White, D. Szöllősi, T. Hussein, K. Kuchler, I.D. Kerr, T. Stockner, J.Y. Lee, Picky ABCG5/G8 and promiscuous ABCG2 - a tale of fatty diets and drug toxicity, FEBS Lett. 594 (2020) 4035–4058. https://doi.org/10.1002/1873-3468.13938.

[24] N. Khunweeraphong, D. Szöllősi, T. Stockner, K. Kuchler, The ABCG2 multidrug transporter is a pump gated by a valve and an extracellular lid, Nat. Commun. 10 (2019) 5433. https://doi.org/10.1038/s41467-019-13302-2.

[25] J.Y. Lee, L.N. Kinch, D.M. Borek, J. Wang, J. Wang, I.L. Urbatsch, X.S. Xie, N. V. Grishin, J.C. Cohen, Z. Otwinowski, H.H. Hobbs, D.M. Rosenbaum, Crystal structure of the human sterol transporter ABCG5/ABCG8, Nature. 533 (2016) 561–564. https://doi.org/10.1038/nature17666.

[26] L. Skarda, J. Kowal, K.P. Locher, Structure of the Human Cholesterol Transporter ABCG1, J. Mol. Biol. 433 (2021) 167218. https://doi.org/https://doi.org/10.1016/j.jmb.2021.167218.

[27] Y. Sun, J. Wang, T. Long, X. Qi, L. Donnelly, N. Elghobashi-Meinhardt, L. Esparza, J.C. Cohen, X.-S. Xie, H.H. Hobbs, X. Li, Molecular basis of cholesterol efflux via ABCG subfamily transporters, Proc. Natl. Acad. Sci. U. S. A. 118 (2021) e2110483118. https://doi.org/10.1073/pnas.2110483118.

[28] M. Ristovski, D. Farhat, S.E.M. Bancud, J.-Y. Lee, Lipid Transporters Beam Signals from Cell Membranes, Membranes (Basel). 11 (2021) 562. https://doi.org/10.3390/membranes11080562.

[29] B.M. Xavier, A.A. Zein, A. Venes, J. Wang, J.Y. Lee, Transmembrane polar relay drives the allosteric regulation for ABCG5/G8 sterol transporter, Int. J. Mol. Sci. 21 (2020) 1–20. https://doi.org/10.3390/ijms21228747.

[30] Á. Telbisz, C. Özvegy-Laczka, T. Hegedűs, A. Váradi, B. Sarkadi, Effects of the lipid environment, cholesterol and bile acids on the function of the purified and reconstituted human ABCG2 protein, Biochem. J. 450 (2013) 387–395. https://doi.org/10.1042/BJ20121485.

[31] G. Gimpl, Interaction of G protein coupled receptors and cholesterol, Chem. Phys. Lipids. (2016) 61–73. https://doi.org/10.1016/j.chemphyslip.2016.04.006.

[32] D. Xu, Y. Li, F. Yang, C.-R. Sun, J. Pan, L. Wang, Z.-P. Chen, S.-C. Fang, X. Yao, W.-T. Hou, C.-Z. Zhou, Y. Chen, Structure and transport mechanism of the human cholesterol transporter ABCG1, Cell Rep. 38 (2022) 110298. https://doi.org/https://doi.org/10.1016/j.celrep.2022.110298.

[33] H. Zhang, C.-S. Huang, X. Yu, J. Lee, A. Vaish, Q. Chen, M. Zhou, Z. Wang, X. Min, Cryo-EM structure of ABCG5/G8 in complex with modulating antibodies, Commun. Biol. 4 (2021) 526. https://doi.org/10.1038/s42003-021-02039-8.

[34] A. Harris, M. Wagner, D. Du, S. Raschka, L.-M. Nentwig, H. Gohlke, S.H.J. Smits, B.F. Luisi, L. Schmitt, Structure and efflux mechanism of the yeast pleiotropic drug resistance transporter Pdr5, Nat. Commun. 12 (2021) 5254. https://doi.org/10.1038/s41467-021-25574-8.

[35] M.J. Ferreiro, C. Pérez, M. Marchesano, S. Ruiz, A. Caputi, P. Aguilera, R. Barrio, R. Cantera, Drosophila melanogaster White Mutant w(1118) Undergo Retinal Degeneration, Front. Neurosci. 11 (2018) 732. https://doi.org/10.3389/fnins.2017.00732.

[36] Y. Mahé, Y. Lemoine, K. Kuchler, The ATP Binding Cassette Transporters Pdr5 and Snq2 of <em>Saccharomyces cerevisiae</em> Can Mediate Transport of Steroids <em>in Vivo</em>*, J. Biol. Chem. 271 (1996) 25167–25172. https://doi.org/10.1074/jbc.271.41.25167.

[37] L. Yu, K. von Bergmann, D. Lutjohann, H.H. Hobbs, J.C. Cohen, Selective sterol accumulation in ABCG5/ABCG8-deficient mice, J. Lipid Res. 45 (2004) 301–307. https://doi.org/10.1194/jlr.M300377-JLR200.

[38] W.J. Allen, T.E. Balius, S. Mukherjee, S.R. Brozell, D.T. Moustakas, P.T. Lang, D.A. Case, I.D. Kuntz, R.C. Rizzo, DOCK 6: Impact of new features and current docking performance, J. Comput. Chem. 36 (2015) 1132–1156. https://doi.org/10.1002/jcc.23905.

[39] B.J. Orlando, M. Liao, ABCG2 transports anticancer drugs via a closed-to-open switch, Nat. Commun. 11 (2020) 2264. https://doi.org/10.1038/s41467-020-16155-2.

[40] S.M. Jackson, I. Manolaridis, J. Kowal, M. Zechner, N.M.I. Taylor, M. Bause, S. Bauer, R. Bartholomaeus, G. Bernhardt, B. Koenig, A. Buschauer, H. Stahlberg, K.-H. Altmann, K.P. Locher, Structural basis of small-molecule inhibition of human multidrug transporter ABCG2, Nat. Struct. Mol. Biol. 25 (2018) 333–340. https://doi.org/10.1038/s41594-018-0049-1.

[41] I. Manolaridis, S.M. Jackson, N.M.I. Taylor, J. Kowal, H. Stahlberg, K.P. Locher, Cryo-EM structures of a human ABCG2 mutant trapped in ATP-bound and substrate-bound states, Nature. 563 (2018) 426–430. https://doi.org/10.1038/s41586-018-0680-3.

[42] D. Sehnal, R. Svobodová Vařeková, K. Berka, L. Pravda, V. Navrátilová, P. Banáš, C.-M. Ionescu, M. Otyepka, J. Koča, MOLE 2.0: advanced approach for analysis of biomacromolecular channels., J. Cheminform. 5 (2013) 39. https://doi.org/10.1186/1758-2946-5-39.

[43] A.I. Greenwood, S. Tristram-Nagle, J.F. Nagle, Partial molecular volumes of lipids and cholesterol., Chem. Phys. Lipids. 143 (2006) 1–10. https://doi.org/10.1016/j.chemphyslip.2006.04.002.

[44] Q. Yu, D. Ni, J. Kowal, I. Manolaridis, S.M. Jackson, H. Stahlberg, K.P. Locher, Structures of ABCG2 under turnover conditions reveal a key step in the drug transport mechanism, Nat. Commun. 12 (2021) 4376. https://doi.org/10.1038/s41467-021-24651-2.

[45] N.M.I. Taylor, I. Manolaridis, S.M. Jackson, J. Kowal, H. Stahlberg, K.P. Locher, Structure of the human multidrug transporter ABCG2, Nature. 546 (2017) 504–509. https://doi.org/10.1038/nature22345.

[46] T. Gose, T. Shafi, Y. Fukuda, S. Das, Y. Wang, A. Allcock, A. Gavan McHarg, J. Lynch, T. Chen, I. Tamai, A. Shelat, R.C. Ford, J.D. Schuetz, ABCG2 requires a single aromatic amino acid to “clamp” substrates and inhibitors into the binding pocket, FASEB J. 34 (2020) 4890–4903. https://doi.org/10.1096/fj.201902338RR.

[47] T. Nagy, Á. Tóth, Á. Telbisz, B. Sarkadi, H. Tordai, A. Tordai, T. Hegedűs, The transport pathway in the ABCG2 protein and its regulation revealed by molecular dynamics simulations, Cell. Mol. Life Sci. 78 (2021) 2329–2339. https://doi.org/10.1007/s00018-020-03651-3.

[48] H. Qian, X. Zhao, P. Cao, J. Lei, N. Yan, X. Gong, Structure of the Human Lipid Exporter ABCA1, Cell. 169 (2017) 1228–1239.e10. https://doi.org/10.1016/j.cell.2017.05.020.

[49] X. Qi, P. Schmiege, E. Coutavas, J. Wang, X. Li, Structures of human Patched and its complex with native palmitoylated sonic hedgehog, Nature. 560 (2018) 128–132. https://doi.org/10.1038/s41586-018-0308-7.

[50] T. Long, X. Qi, A. Hassan, Q. Liang, J.K. De Brabander, X. Li, Structural basis for itraconazole-mediated NPC1 inhibition, Nat. Commun. 11 (2020) 152. https://doi.org/10.1038/s41467-019-13917-5.

[51] J. Jumper, R. Evans, A. Pritzel, T. Green, M. Figurnov, O. Ronneberger, K. Tunyasuvunakool, R. Bates, A. Žídek, A. Potapenko, A. Bridgland, C. Meyer, S.A.A. Kohl, A.J. Ballard, A. Cowie, B. Romera-Paredes, S. Nikolov, R. Jain, J. Adler, T. Back, S. Petersen, D. Reiman, E. Clancy, M. Zielinski, M. Steinegger, M. Pacholska, T. Berghammer, S. Bodenstein, D. Silver, O. Vinyals, A.W. Senior, K. Kavukcuoglu, P. Kohli, D. Hassabis, Highly accurate protein structure prediction with AlphaFold, Nature. 596 (2021) 583–589. https://doi.org/10.1038/s41586-021-03819-2.

[52] O. Sano, A. Kobayashi, K. Nagao, K. Kumagai, N. Kioka, K. Hanada, K. Ueda, M. Matsuo, Sphingomyelin-dependence of cholesterol efflux mediated by ABCG1, J. Lipid Res. 48 (2007) 2377–2384. https://doi.org/10.1194/jlr.M700139-JLR200.

[53] I. Ermilova, A.P. Lyubartsev, Cholesterol in phospholipid bilayers: positions and orientations inside membranes with different unsaturation degrees, Soft Matter. 15 (2019) 78–93. https://doi.org/10.1039/C8SM01937A.

[54] Z. Otwinowski, W.B.T.-M. in E. Minor, [20] Processing of X-ray diffraction data collected in oscillation mode, in: Macromol. Crystallogr. Part A, Academic Press, 1997: pp. 307–326. https://doi.org/https://doi.org/10.1016/S0076-6879(97)76066-X.

[55] M.J. Grayling, PhaseR: An R package for phase plane analysis of autonomous ODE systems, R J. 6 (2014) 43–51. https://doi.org/10.32614/rj-2014-023.

[56] P.D. Adams, P. V Afonine, G. Bunkóczi, V.B. Chen, I.W. Davis, N. Echols, J.J. Headd, L.-W. Hung, G.J. Kapral, R.W. Grosse-Kunstleve, A.J. McCoy, N.W. Moriarty, R. Oeffner, R.J. Read, D.C. Richardson, J.S. Richardson, T.C. Terwilliger, P.H. Zwart, PHENIX: a comprehensive Python-based system for macromolecular structure solution, Acta Crystallogr. D. Biol. Crystallogr. 66 (2010) 213–221. https://doi.org/10.1107/S0907444909052925.

[57] P. Emsley, K. Cowtan, {\it Coot}: model-building tools for molecular graphics, Acta Crystallogr. Sect. D. 60 (2004) 2126–2132. https://doi.org/10.1107/S0907444904019158.

[58] V.B. Chen, W.B. Arendall 3rd, J.J. Headd, D.A. Keedy, R.M. Immormino, G.J. Kapral, L.W. Murray, J.S. Richardson, D.C. Richardson, MolProbity: all-atom structure validation for macromolecular crystallography, Acta Crystallogr. D. Biol. Crystallogr. 66 (2010) 12– 21. https://doi.org/10.1107/S0907444909042073.

[59] P.J.A. Cock, T. Antao, J.T. Chang, B.A. Chapman, C.J. Cox, A. Dalke, I. Friedberg, T. Hamelryck, F. Kauff, B. Wilczynski, M.J.L. de Hoon, Biopython: freely available Python tools for computational molecular biology and bioinformatics, Bioinformatics. 25 (2009) 1422–1423. https://doi.org/10.1093/bioinformatics/btp163.

[60] S. Henikoff, J.G. Henikoff, Amino acid substitution matrices from protein blocks, Proc. Natl. Acad. Sci. U. S. A. 89 (1992) 10915–10919.b https://doi.org/10.1073/pnas.89.22.10915.

[61] J.D. Thompson, D.G. Higgins, T.J. Gibson, CLUSTAL W: improving the sensitivity of progressive multiple sequence alignment through sequence weighting, position-specific gap penalties and weight matrix choice, Nucleic Acids Res. 22 (1994) 4673–4680. https://doi.org/10.1093/nar/22.22.4673.

[62] J. Pei, B.-H. Kim, N. V Grishin, PROMALS3D: a tool for multiple protein sequence and structure alignments, Nucleic Acids Res. 36 (2008) 2295–2300. https://doi.org/10.1093/nar/gkn072.

[63] R.C. Edgar, MUSCLE: multiple sequence alignment with high accuracy and high throughput, Nucleic Acids Res. 32 (2004) 1792–1797. https://doi.org/10.1093/nar/gkh340.

[64] C.D. Livingstone, G.J. Barton, Protein sequence alignments: a strategy for the hierarchical analysis of residue conservation, Bioinformatics. 9 (1993) 745–756. https://doi.org/10.1093/bioinformatics/9.6.745.

[65] B. Webb, A. Sali, Protein Structure Modeling with MODELLER BT - Protein Structure Prediction, in: D. Kihara (Ed.), Springer New York, New York, NY, 2014: pp. 1–15. https://doi.org/10.1007/978-1-4939-0366-5_1.

[66] C.J. Williams, J.J. Headd, N.W. Moriarty, M.G. Prisant, L.L. Videau, L.N. Deis, V. Verma, D.A. Keedy, B.J. Hintze, V.B. Chen, S. Jain, S.M. Lewis, W.B. Arendall 3rd, J. Snoeyink, P.D. Adams, S.C. Lovell, J.S. Richardson, D.C. Richardson, MolProbity: More and better reference data for improved all-atom structure validation, Protein Sci. 27 (2018) 293–315. https://doi.org/10.1002/pro.3330.

[67] M. V Shapovalov, R.L. Dunbrack Jr, A smoothed backbone-dependent rotamer library for proteins derived from adaptive kernel density estimates and regressions, Structure. 19 (2011) 844–858. https://doi.org/10.1016/j.str.2011.03.019.

[68] J. Wang, W. Wang, P.A. Kollman, D.A. Case, Automatic atom type and bond type perception in molecular mechanical calculations, J. Mol. Graph. Model. 25 (2006) 247– 260. https://doi.org/https://doi.org/10.1016/j.jmgm.2005.12.005.

[69] A. Morin, B. Eisenbraun, J. Key, P.C. Sanschagrin, M.A. Timony, M. Ottaviano, P. Sliz, Collaboration gets the most out of software, Elife. 2 (2013) e01456. https://doi.org/10.7554/eLife.01456.

[70] E.F. Pettersen, T.D. Goddard, C.C. Huang, G.S. Couch, D.M. Greenblatt, E.C. Meng, T.E. Ferrin, UCSF Chimera - A visualization system for exploratory research and analysis, J. Comput. Chem. 25 (2004) 1605–1612. https://doi.org/10.1002/jcc.20084.

[71] A. Radhakrishnan, L.-P. Sun, H.J. Kwon, M.S. Brown, J.L. Goldstein, Direct Binding of Cholesterol to the Purified Membrane Region of SCAP: Mechanism for a Sterol-Sensing Domain, Mol. Cell. 15 (2004) 259–268. https://doi.org/10.1016/j.molcel.2004.06.019.

